# Machine Learning-Enhanced Drug Discovery for BACE1: A Novel Approach to Alzheimer’s Therapeutics

**DOI:** 10.1101/2024.09.24.614844

**Authors:** Satyam Sangeet

## Abstract

Alzheimer’s disease (AD) remains a significant global health challenge, with BACE1, a critical therapeutic target due to its role in the amyloidogenic pathway. This study presents an innovative approach to drug development by integrating machine learning (ML) techniques to identify the potential BACE1 inhibitors. Utilizing a comprehensive dataset of protein-ligand interactions, the current study employed a machine learning model to generate novel ligands with high binding affinity and specificity for the BACE1 active site. The model’s efficacy was validated through molecular docking studies, which demonstrated superior binding affinities compared to existing FDA-approved inhibitors such as Atabecestat and Lanabecestat. The findings reveal that the top candidate, MLC10, exhibits a unique mechanism of action by promoting controlled flexibility in BACE1, contrasting with the rigid conformations induced by traditional inhibitors. This dynamic modulation enhances the enzyme’s inhibition, suggesting a promising avenue for therapeutic intervention. Furthermore, solvent-accessible surface area analyses indicate that MLC10 facilitates a more favourable protein conformation, potentially altering catalytic behaviour through increased solvent interactions. This work underscores the transformative potential of machine learning in drug discovery, paving the way for the development of next-generation BACE1 inhibitors. By harnessing computational techniques, the present work aims to address the limitations of current therapies and contribute to more effective treatments for Alzheimer’s disease, ultimately improving patient outcomes and quality of life.

## Introduction

Alzheimer’s disease stands at the forefront of neurodegenerative disorders, presenting an unparalleled challenge to modern medicine and an ever-increasing socioeconomic burden worldwide [1, 2]. At the molecular nexus of AD pathology lies β-site amyloid precursor protein cleaving enzyme 1 (BACE1), a critical mediator in the amyloidogenic pathway responsible for generating neurotoxic amyloid-beta (Aβ) peptides. These peptides aggregate to form the characteristic amyloid plaques that, along with neurofibrillary tangles, define the histopathological hallmarks of AD [3, 4, 5]. The centrality of BACE1 in AD pathogenesis [6, 7, 8, 9] has positioned it as a prime target for therapeutic intervention, with the strategic aim of attenuating Aβ production. BACE1’s catalytic machinery involve two aspartate residues, Asp32 and Asp228 [10, 11] (Fig 1, surface view, yellow), which coordinate a water molecule to facilitate peptide bond hydrolysis [12]. The complexity of BACE1 extends beyond its catalytic core, encompassing dynamic structural elements that modulate its activity. Most notable is the flap region (residues 67-77) (Fig 1, cartoon, red), a flexible hairpin loop above the active site cleft. This flap transitions from an open state to a closed conformation upon substrate or inhibitor binding, crucial for stabilizing the enzyme-substrate complex [8, 9]. Additionally, the 10s loop (residues 5-16, Fig 1, cartoon view, green) undergoes significant conformational changes, regulating substrate access to the active site [13]. The interplay between these structural components creates a sophisticated molecular mechanism finely tuned for its physiological function. This structural complexity not only influences BACE1’s substrate specificity but also presents unique challenges and opportunities in inhibitor design. Despite the compelling rationale and promising preclinical data supporting BACE1 inhibition as a therapeutic strategy, the translation of these findings into clinically efficacious treatments has proven remarkably challenging. A series of clinical trials investigating BACE1 inhibitors, including Atabecestat [14], Lanabecestat [15, 16], and Verubecestat [17], have yielded disappointing results, failing to demonstrate significant cognitive benefits in AD patients. These setbacks have not only highlighted the complexities inherent in targeting the amyloid pathway but have also underscored the limitations of traditional drug development paradigms in addressing the multifaceted nature of AD pathophysiology.

**Fig. 1:**
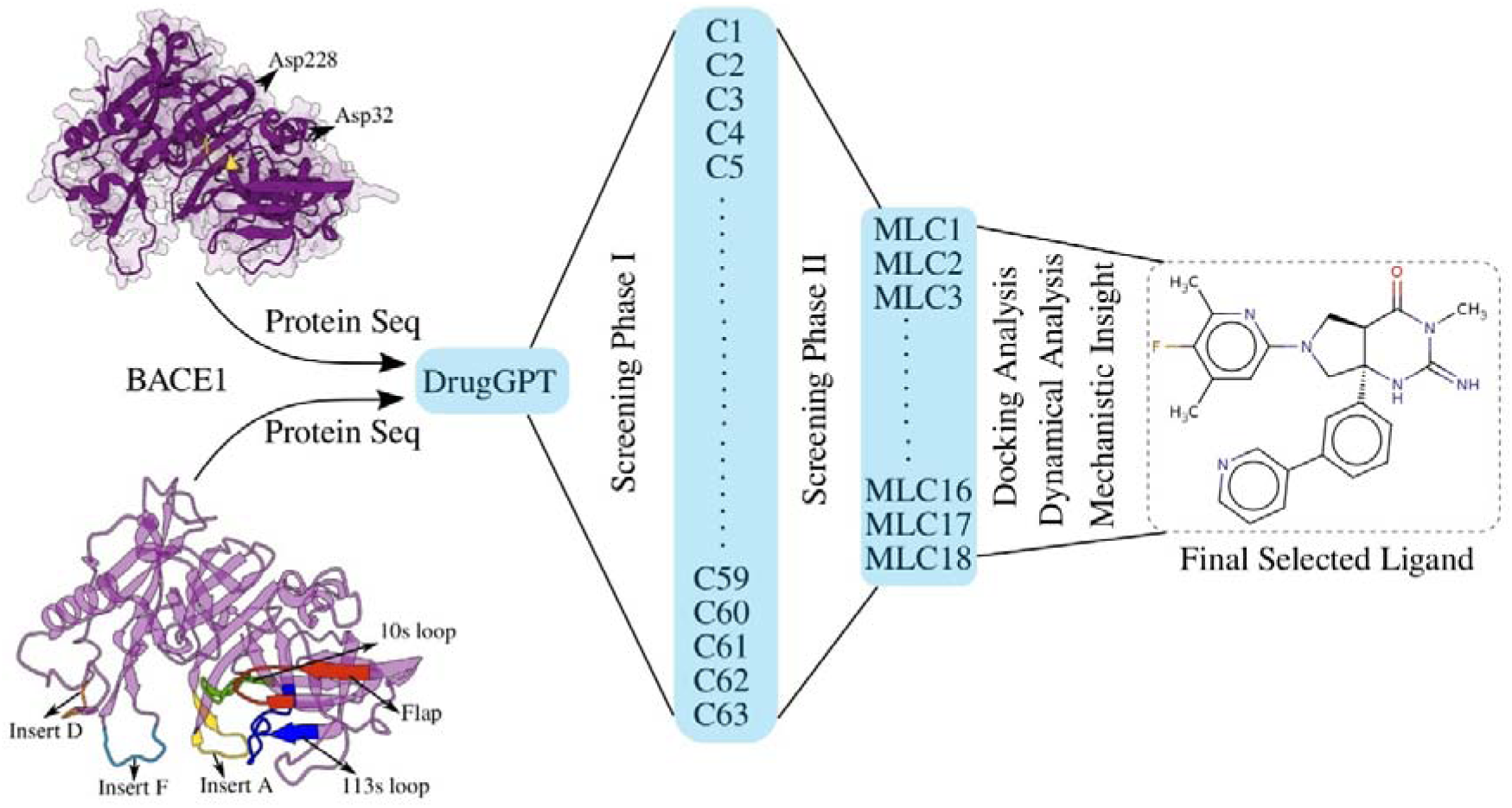
Overview of the study integrating ML with classical drug development pipeline. The surface and cartoon view of the BACE1 structure representing active site Aspartates and the important regions involved in the substrate binding events

The recurrent failures in AD drug development necessitate a shift in our approach to therapeutic discovery and optimization. In this context, machine learning (ML) has emerged as a transformative force, offering unprecedented capabilities to navigate the vast chemical space [18, 19, 20], decipher complex biological interactions [21, 22], and predict therapeutic outcomes with increasing accuracy [23, 24]. By leveraging advanced algorithms and computational models, ML technologies can process and analyse diverse datasets at scales far beyond human capacity, enabling the identification of subtle patterns and relationships that may elude conventional analytical methods. The integration of ML into the drug discovery pipeline presents a multitude of advantages that address many of the bottlenecks encountered in traditional approaches. This study represents a concerted effort to harness the power of ML in the pursuit of novel, efficacious BACE1 inhibitors for the treatment of AD. The current study utilises DrugGPT [25], a generative model tailored for drug discovery, to explore chemical space and design novel ligands. DrugGPT integrates a sophisticated tokenization strategy that decomposes both protein sequences and ligands into discrete lexical units, enabling it to model complex chemical and biological interactions. By employing this strategy, the model can efficiently traverse vast chemical spaces to generate molecules that possess desirable therapeutic properties. The model presented potential molecules that may poses the ability as potent BACE1 inhibitors. Through this approach, the present study identified one promising lead compound, MLC10, as an effective BACE1 inhibitor. The results presented through this study has the potential to inform future drug development efforts across a range of neurodegenerative disorders, contributing to a broader understanding of how ML can be leveraged to address complex biological challenges.

## Methodology

### Ligand Generation using DrugGPT

This methodology leverages DrugGPT’s [25] ability to explore chemical space in a targeted, context-aware manner, ensuring that the generated ligands are not only chemically diverse but also relevant to the biological target. By utilizing the BACE1 protein sequence as input, the study ensured that the generated ligands are structurally predisposed to interact with BACE1’s active site. This enhances the likelihood of identifying novel molecules that could serve as effective BACE1 inhibitors, a key therapeutic strategy for mitigating Alzheimer’s disease progression. The model was deployed locally resulting in the generation of SMILES notation of ligands.

### Initial Screening

Following the generation of ligand SMILES by the model, corresponding 2D molecular structures were created and subjected to a two-stage initial screening process. In the first phase, Lipinski’s rule of five was rigorously applied using the SIMANA server [26]. A stringent criterion was enforced, whereby any ligand exhibiting even a single violation of the rule was excluded from further analysis. In the second phase, the LD50 value for each ligand was calculated, using ProTox 3 server [27], with a threshold of ≥2000 mg/kg employed to ensure a favourable safety profile. Ligands falling below this toxicity cutoff were similarly discarded from further consideration.

### Molecular Docking

To assess the inhibitory potential of novel ligands generated via DrugGPT against BACE1, the study employed sophisticated molecular docking simulations using AutoDock Vina [28]. The high-resolution crystal structure of BACE1 (PDB ID: 2OHM) was retrieved from the Protein Data Bank [29] and meticulously prepared for in silico analysis. This preparation began with the removal of extraneous elements, including small ionic species and water molecules, which could potentially interfere with the docking simulations. Subsequently, the structure was subjected to energy minimization using the Chimera software package [30]. The energy minimization protocol employed a two-stage approach. Initially, the Steepest Descent algorithm was applied for 1000 iterations, using a step size of 0.1 Å. This was followed by 100 steps of conjugate gradient minimization, also with a 0.1 Å step size. This dual-method approach ensures a thorough exploration of the conformational space, resulting in an energetically favourable protein structure [31]. For the force field assignments, AMBER ff14SB force field [32] was used. Non-standard residues were parameterized using the AMI-BCC force field. MolStar [33] was used for visualisation. Furthermore, the ligands were energy minimised using the MM2 energy minimisation algorithm [34] implemented in Chem3D [35]. Detailed results from each docking procedure are available in Supplementary Table S1. To validate the docking protocol, redocking was performed using N3-benzylpyridine-2,3-diamine (8AP), the co-crystallized inhibitor present in the 2OHM PDB structure. Validation was quantitatively assessed by calculating the Root Mean Square Deviation (RMSD) between the docked pose and the original crystal-bound conformation of 8AP.

### Molecular Dynamic Simulation

Molecular dynamics (MD) simulations were conducted to investigate the dynamic interactions of known inhibitors and top-ranked phytochemicals with BACE1. Each simulation was performed in triplicate using GROMACS version 2018.1 [36, 37, 38], running for 150 nanoseconds (ns). The protein-ligand complexes were prepared with the CHARMM36 force field [39], and ligand topologies were generated using CGenFF [40]. The system was solvated in a 5 nm X 5nm X 5nm cubic box with TIP3P water molecules, and charge neutrality was achieved by adding Na□ and Cl□ ions. Energy minimization was carried out using the steepest descent algorithm for up to 5000 steps. The system was then equilibrated in two stages: first under an NVT ensemble at 300 K with a V-rescale thermostats for 1 ns, followed by an NPT ensemble at 1 bar pressure using a Berendsen barostat for another 1 ns. The production MD phase ran for 150 ns, employing the leap-frog algorithm with a 2-fs time step. Van der Waals interactions were treated with a 1.2 nm cutoff, while long-range electrostatic interactions were computed using the Particle Mesh Ewald (PME) method. Temperature was controlled with the Nosé-Hoover thermostat, and pressure was maintained using the Parrinello-Rahman barostat. System coordinates were saved every 20 ps for subsequent analysis, which focused on assessing the stability, flexibility, and conformational behaviour of the protein-ligand complexes throughout the simulation.

### Molecular Mechanics Poisson Boltzmann Surface Area (MMPBSA)

The binding free energy (Δ*G_bind_*) between a protein and its ligand is a critical parameter for elucidating molecular interactions within protein-ligand complexes. In this study, we employed the Molecular Mechanics/Poisson-Boltzmann Surface Area (MM/PBSA) method, a well-established approach for estimating binding free energy. The binding free energy (Δ*G_bind_*) is calculated as the difference between the free energy of the protein-ligand complex (Δ*G_complex_*) and the sum of the free energies of the isolated protein (Δ*G_protein_*) and ligand (Δ*G_ligand_*), expressed as:

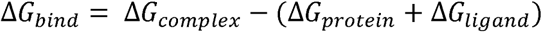

Here, Δ*G_complex_* represents the free energy of the protein-ligand complex, while Δ*G_protein_* and Δ*G_ligand_* correspond to the free energies of the unbound protein and ligand, respectively. This approach offers a comprehensive energetic framework for evaluating the stability and binding affinity of protein-ligand interactions.

### Dynamic Cross Correlation Matrix (DCCM) Analysis

Dynamic cross-correlation analysis (DCCM) was performed, using SIMANA [26], to investigate the relationships between protein residues. A structural ensemble was created from the MD simulation trajectories of the protein-ligand complexes with the highest-ranked ligands. The correlations between residues were represented in a matrix, visualized as a Dynamic Cross-Correlation Matrix (DCCM). To ensure the analysis was robust, three independent MD simulation trajectories were run, the results were averaged, and the final DCCM was calculated based on these combined data [41].

### Principal Component Analysis (PCA)

Principal Component Analysis (PCA) was performed to understand the structural dynamics of the protein-ligand complex, expressed in thermal energy (*k_B_T*). The results were analysed by plotting the first principal component (PC1) against the second (PC2), with each data point representing a distinct conformational state of the complex. To do this, covariance matrix was computed, which shows how variables relate to each other and how changes in one variable affect others. Then eigen decomposition of the covariance matrix was performed, obtaining eigenvalues and eigenvectors. The eigenvalues indicate the amount of variance explained by each principal component, while the eigenvectors represent the directions of these components in the original feature space. A projection matrix was created to map the original data into a lower-dimensional space, which highlighted the significant principal components and their role in the protein-ligand complex’s dynamics. Finally, the loadings of the original variables were analysed on each principal component to determine which interactions and measurements most influenced the variance captured by each component thereby representing the “essential dynamics” of the protein.

### Deformity and Fluctuation Analysis

To evaluate the deformability, structural stability and energetic fluctuation of BACE1 in the presence of inhibitors and generated ligand, Normal Mode Analysis (NMA) was performed via the iMOD server [42]. The equilibrated structures from the MD simulations were provided to the server. NMA involved creating an elastic network model of the protein, where each Cα atom is modelled as a node connected by springs representing interatomic interactions. The server then calculated the collective motions of the protein along the lowest-frequency normal modes. Deformability was analysed by determining the deformation energy for each residue, which reflects how much each residue can change conformation. Regions with high deformability are more flexible, while those with low deformability are more rigid. Moreover, fluctuation analysis was also performed reflecting the residual fluctuations of individual atoms or residues due to the normal modes of vibration, representing how much each residue moves in response to different vibrational modes. With respect to NMA, these fluctuations relate to the energy landscape and indicate how flexible each residue is within the low-frequency vibrational modes. Thus, the fluctuation values depict the energy fluctuations derived from the NMA modes.

## Results

### Drug Selection & Initial Screening

The current study employed an innovative approach integrating machine learning (ML) with traditional drug discovery methods to identify potential BACE1 inhibitors. DrugGPT model was utilized to generate ligands targeting BACE1. By utilizing the amino acid sequence of BACE1, the goal was to generate novel ligands with high binding affinity and specificity for this protein. The model was trained on an extensive dataset of protein-ligand interactions [25], equipping it to understand the complex relationships between protein structures and their respective ligands. After inputting the BACE1 protein sequence into the DrugGPT model, 63 unique SMILES notations (initially labelled C1 through C63, S1 Table), each representing a distinct chemical structure was generated. To evaluate the quality and diversity of these generated ligands, an in-depth analysis of the SMILES notations was performed. The ligands exhibited a broad range of structural characteristics (S1 Fig), including differences in functional groups, ring structures, and molecular weights. Notably, the model successfully generated ligands with varying levels of complexity (S2 Fig), which is crucial for optimizing binding interactions with the BACE1 active site (S1 Fig). These compounds were subsequently subjected to a rigorous, multi-step selection process. First, all 63 compounds underwent evaluation using Lipinski’s rule of five, via SIMANA server [26], to assess their drug-likeness and potential for oral bioavailability (S2 Table). This initial screening reduced the compound pool to 18 candidates (renamed MLC1 through MLC18).

The remaining 18 compounds were then evaluated for acute toxicity using LD50 predictions (Table 1). This toxicity screening further narrowed our selection to two promising compounds: MLC10 and MLC11 (Fig 3). The potential efficacy of these two selected compounds were assessed using molecular docking analyses. As controls, we included three established BACE1 inhibitors: Atabecestat, Lanabecestat, and Verubecestat (Fig 2).

**Fig 2.**
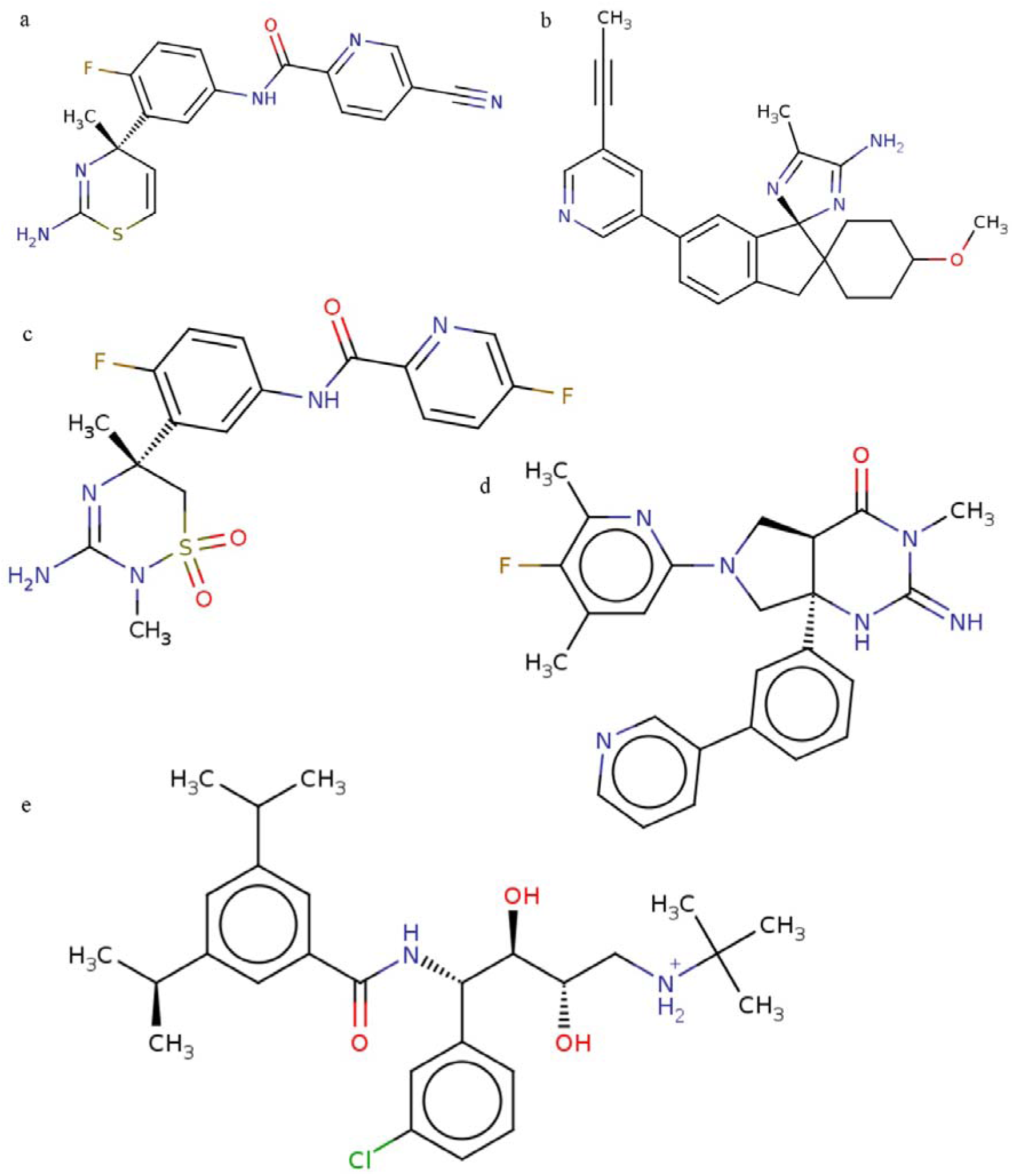
2D structure of known Inhibitors and ML generated top hit ligands (a) Atabecestat (b) Lanabecestat (c) Verubecestat (d) MLC10 (e) MLC11

**Fig 3:**
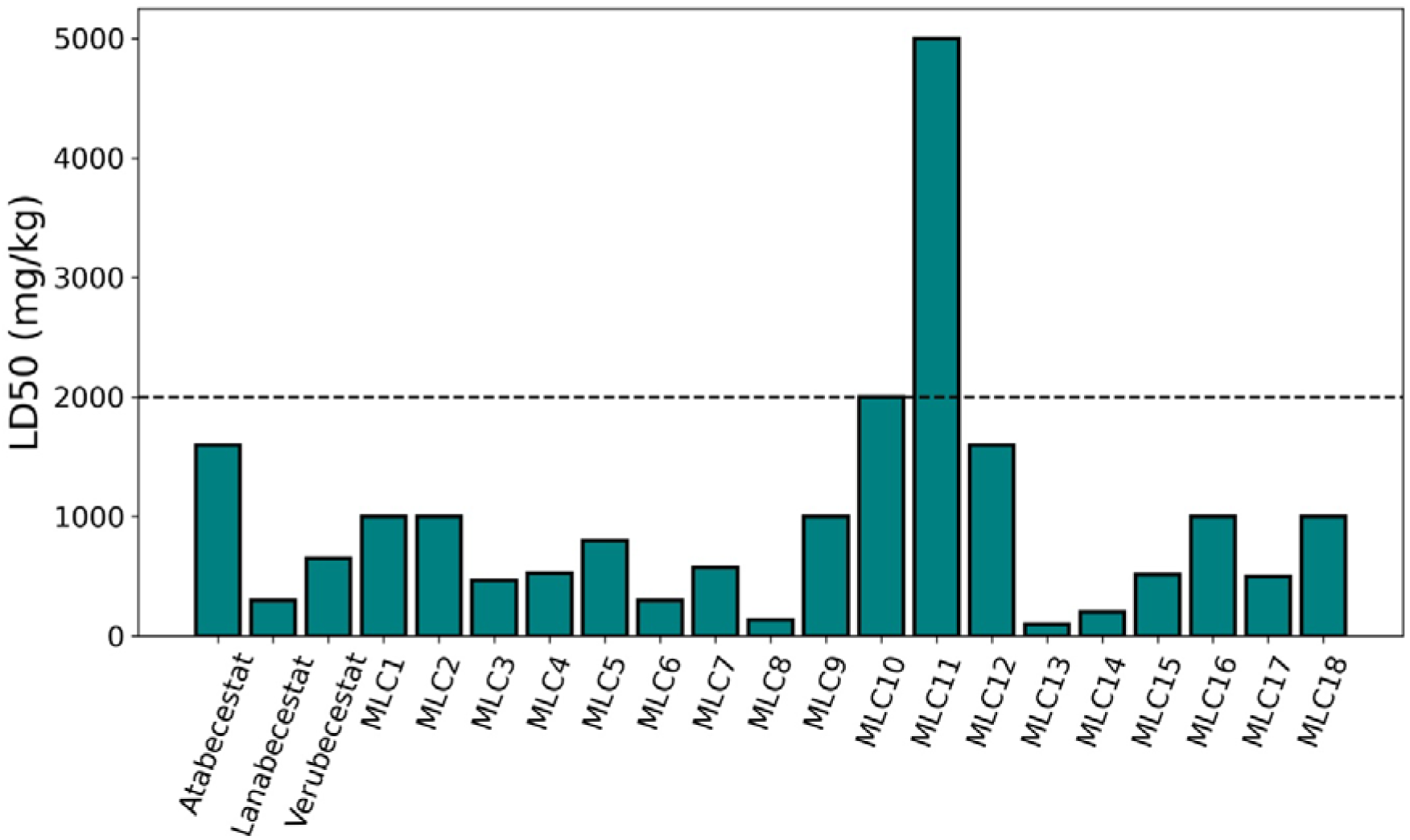
LD50 values of known inhibitors and the top 18 generated molecules. The cutoff (dashed line) is set at >= 2000 mg/kg. MLC10 and MLC11 showed a LD50 value of 2000 mg/kg and 5000 mg/kg respectively

**Table 1:** Initial screening of known inhibitors and generated ligands.

| Compound | Lipinski Rule* |  |  |  | LD50 (mg/kg) |
| --- | --- | --- | --- | --- | --- |
|  | R1 | R2 | R3 | R4 |  |
| Atabecestat | 367.41 | 2 | 6 | 3.14 | 1600 |
| Lanabecestat | 412.54 | 1 | 5 | 4.24 | 300 |
| Verubecestat | 409.42 | 2 | 6 | 1.42 | 650 |
| MLC1 | 463.62 | 5 | 5 | 1.84 | 1000 |
| MLC2 | 449.64 | 3 | 5 | 1.78 | 1000 |
| MLC3 | 458.59 | 3 | 8 | 1.80 | 465 |
| MLC4 | 409.43 | 0 | 5 | 2.54 | 525 |
| MLC5 | 415.44 | 2 | 4 | 2.13 | 800 |
| MLC6 | 446.54 | 2 | 3 | 4.26 | 300 |
| MLC7 | 479.60 | 3 | 3 | 3.88 | 572 |
| MLC8 | 477.52 | 3 | 8 | 1.64 | 135 |
| MLC9 | 434.41 | 3 | 4 | 3.86 | 1000 |
| MLC10 | 444.51 | 2 | 5 | 3.23 | 2000 |
| MLC11 | 476.08 | 4 | 3 | 4.14 | 5000 |
| MLC12 | 432.43 | 3 | 5 | 2.40 | 1600 |
| MLC13 | 314.38 | 1 | 2 | 3.75 | 100 |
| MLC14 | 488.57 | 3 | 7 | 3.26 | 200 |
| MLC15 | 445.47 | 0 | 4 | 4.38 | 515 |
| MLC16 | 375.35 | 1 | 5 | 3.48 | 1000 |
| MLC17 | 474.51 | 4 | 10 | 3.25 | 500 |
| MLC18 | 381.44 | 1 | 5 | 2.38 | 1000 |

*\*R1 to R4 denote the Lipinski's RO5 where R1 = Molecular Weight, R2 = Number of Hydrogen bond donors, R3 = Number of Hydrogen bond acceptors and R4 = LogP*

### Molecular Docking

Molecular docking analyses revealed detailed binding interactions between BACE1 and the novel compounds MLC10 and MLC11 (Fig 2), alongside known BACE1 inhibitors Atabecestat, Lanabecestat, and Verubecestat. The binding energies for these compounds were found to be −7.5 kcal/mol for Atabecestat, −8.8 kcal/mol for Lanabecestat, −7.4 kcal/mol for Verubecestat, −8.8 kcal/mol for MLC10, and −7.7 kcal/mol for MLC11. Notably, MLC10 exhibited a binding energy identical to Lanabecestat, the most potent reference inhibitor, suggesting a comparable affinity for the BACE1 active site. Detailed hydrogen bonding analyses (Table 2) showed that MLC10 interacted with critical residues in the BACE1 catalytic site, including Tyr71, Gly74, and Asp228 (Fig 4d). Importantly, its engagement with Asp228, one of the key catalytic dyad residues, supports its potential as an effective BACE1 inhibitor. MLC10 also demonstrated a balance of hydrophobic and electrostatic interactions, forming salt bridges with both catalytic residues Asp32 and Asp228, further reinforcing its inhibitory potential. MLC11, while exhibiting a moderate binding energy (−7.7 kcal/mol), formed an extensive network of hydrophobic interactions with eight key residues, including Leu30, Tyr71, Phe108, and Trp115 (S3 Fig). This broad interaction profile spanned critical regions of BACE1, including the flap region (residues 67-75) and substrate binding site, potentially accounting for its binding efficiency despite fewer hydrogen bonds compared to MLC10. The comprehensive interaction profiles of MLC10 and MLC11 suggest that these compounds effectively engage multiple functional regions of BACE1, including the catalytic dyad, flap region, and substrate binding site. The direct engagement of Asp32 and Asp228 by both compounds indicates potential interference with the catalytic mechanism of BACE1. Furthermore, interactions with residues such as Tyr71 (flap region) and Arg128 (113s loop) suggest a multi-region targeting strategy that could enhance both specificity and potency. These findings provide strong evidence for the inhibitory potential of MLC10 and MLC11, with binding energies and interaction profiles comparable to or exceeding established BACE1 inhibitors. Based on the binding affinity, MLC10 was selected for further investigations.

**Fig 4:**
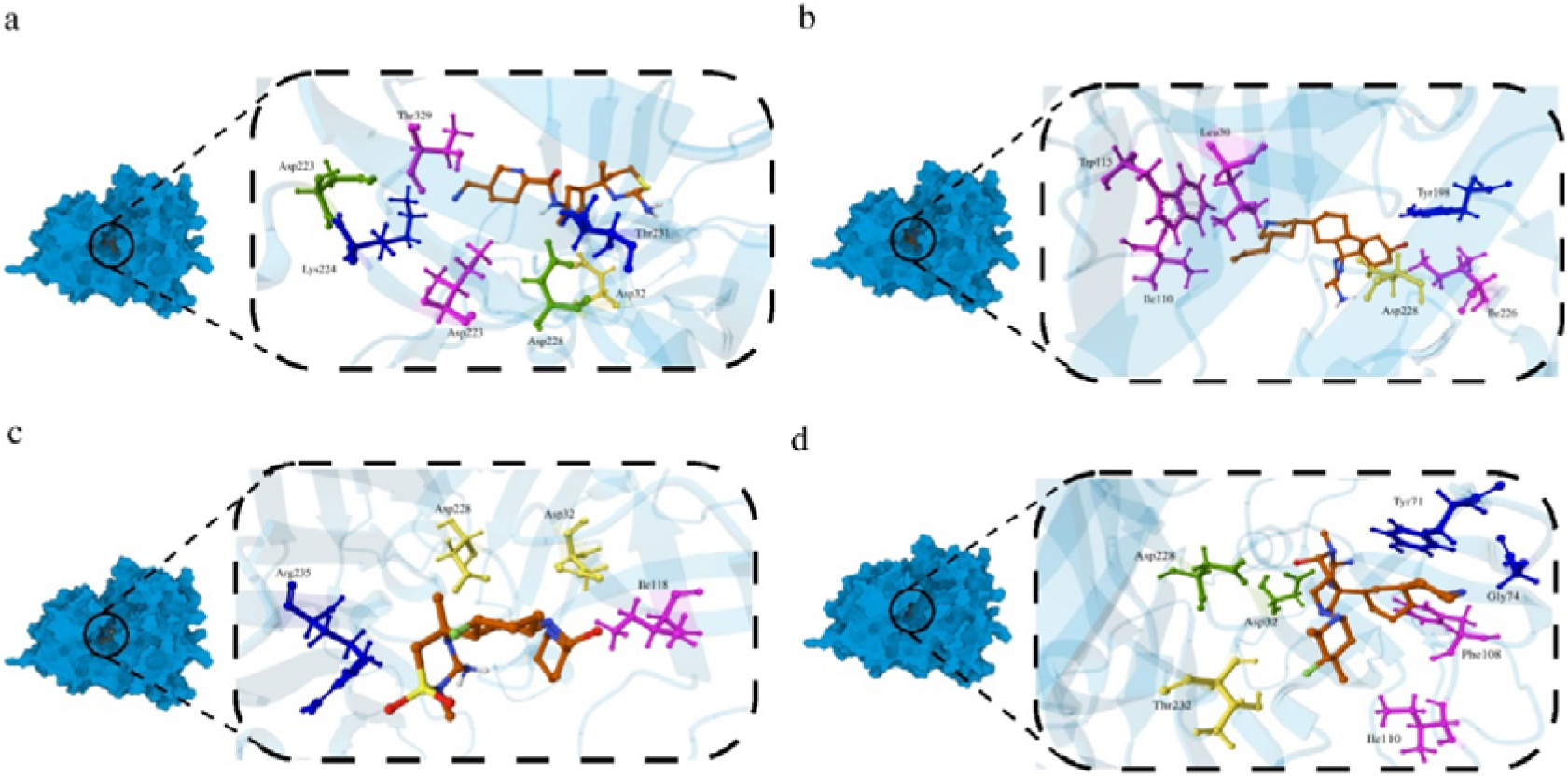
Molecular Docking interaction profile of the BACE1 with known inhibitors and the top hit molecule. (a) Atabecestat (b) Lanabecestat (c) Verubecestat (d) MLC10. Hydrogen Bonding (blue), Hydrophobic interaction (magenta), Salt Bridge (green), Halogen (yellow)

**Table 2:** Hydrogen Bonding exhibited by the known ligands and the top hit generated molecules.

| Ligand | Residue | Distance H-A | Distance D-A | Donor Angle |
| --- | --- | --- | --- | --- |
| Atabecestat | Lys224 | 2.67 | 3.14 | 108.72 |
|  | Thr231 | 2.62 | 3.12 | 110.92 |
| Lanabecestat | Tyr198 | 2.19 | 3.04 | 146.16 |
|  | Asp228 | 3.02 | 3.79 | 133.17 |
| Verubecestat | Asp32 | 2.65 | 3.35 | 129.63 |
|  | Arg235 | 2.77 | 3.12 | 101.18 |
| MLC10 | Tyr71 | 2.52 | 3.02 | 109.92 |
|  | Gly74 | 2.52 | 3.27 | 130.89 |
|  | Asp228 | 2.61 | 3.37 | 135.85 |
| MLC11 | Asp32 | 2.97 | 3.42 | 109.68 |
|  | Gly34 | 3.54 | 4.02 | 110.75 |

**Table 3:** Other interactions apart from hydrogen bonding exhibited by the known inhibitors and the top hit generated molecules.

| Ligand | Residue | Interaction Type | Distance |
| --- | --- | --- | --- |
| Atabecestat | Ile226 | Hydrophobic | 3.66 |
|  | Thr329 | Hydrophobic | 3.87 |
|  | Asp32 | Halogen | 3.83 |
|  | Asp223 | Salt Bridge | 5.39 |
|  | Asp232 | Salt Bridge | 3.31 |
| Lanabecestat | Leu30 | Hydrophobic | 3.70 |
|  | Ile110 | Hydrophobic | 3.94 |
|  | Trp115 | Hydrophobic | 3.83 |
|  | Ile226 | Hydrophobic | 3.98 |
|  | Asp228 | Salt Bridge | 5.44 |
| Verubecestat | Ile118 | Hydrophobic | 3.34 |
|  | Asp32 | Salt Bridge | 3.54 |
|  | Asp228 | Salt Bridge | 5.18 |
| MLC10 | Tyr71 | Hydrophobic | 3.50 |
|  | Phe108 | Hydrophobic | 3.77 |
|  | Ile110 | Hydrophobic | 3.80 |
|  | Thr232 | Halogen | 3.09 |
|  | Asp32 | Salt Bridge | 4.65 |
|  | Asp228 | Salt Bridge | 5.27 |
| MLC11 | Leu30 | Hydrophobic | 3.63 |
|  | Tyr71 | Hydrophobic | 3.77 |
|  | Phe108 | Hydrophobic | 3.97 |
|  | Ile110 | Hydrophobic | 3.59 |
|  | Trp115 | Hydrophobic | 3.60 |
|  | Tyr198 | Hydrophobic | 3.66 |
|  | Thr329 | Hydrophobic | 3.79 |
|  | Val332 | Hydrophobic | 3.64 |

### Redocking Analysis

A comprehensive docking study was performed to assess the interaction between 8AP and the crystalline structure of BACE1 to validate the docking procedure used. The results demonstrated that the docked 8AP exhibited an affinity for the catalytic site of BACE1, with a binding energy of −5.8 kcal/mol. The three-dimensional positioning of the docked 8AP was superimposed with the 8AP ligand in the native crystal structure (S4 Fig), resulting in an RMSD (Root Mean Square Deviation) of 1.1 Å. This alignment confirmed the accuracy and reliability of the docking protocol employed.

### Molecular Dynamic Simulation

#### Root Mean Square Deviation (RMSD)

The 150 ns molecular dynamics simulations revealed significant conformational dynamics of BACE1 in complex with MLC10 and established inhibitors offering insights into their potential mechanisms of action and efficacy. RMSD analyses showed distinct patterns of protein stability and flexibility across different ligand-bound states. The protein-only simulation (black) stabilized at an RMSD value of ∼0.2 nm after equilibration, reflecting the baseline conformational flexibility of BACE1 in the absence of ligands (Fig 5a, black). Among the known inhibitors, Verubecestat (Fig 5a, blue) exhibited a consistent RMSD profile, closely mirroring the protein-only simulation, suggesting that it stabilizes BACE1 in a near-native conformation, potentially contributing to its known efficacy. Atabecestat (Fig 5a, red) and Lanabecestat (Fig 5a, green) displayed more variable RMSD patterns, with Lanabecestat showing notable fluctuations, particularly in the 30-50 ns range, indicating possible induced-fit mechanisms or dynamic binding modes that transiently alter BACE1’s conformation. Notably, MLC10 (Fig 5a, magenta) exhibited a unique RMSD profile, maintaining variable yet close proximity to the protein, indicating a balance between stabilizing BACE1’s conformation and allowing for induced flexibility. This dynamic interaction suggests that MLC10 may optimize binding to key catalytic residues while preserving necessary protein dynamics. The flexibility observed with MLC10 may enhance its inhibitory efficacy through multiple mechanisms. MLC10’s dynamic behaviour could enable induced-fit optimization, allowing the active site geometry to be fine-tuned, thereby maximizing complementarity between the inhibitor and BACE1’s active site. Moreover, MLC10’s flexibility could interact with the BACE1 flap region (residues 67-75), a critical area for substrate access and product release, potentially modulating BACE1 activity and enhancing inhibition. Moreover, MLC10’s RMSD distribution is similar to Verubecestat’s with a bimodal distribution (Fig 5a, distribution curve), yet offering some nuanced differences. The ability of MLC10 to maintain or selectively enhance protein flexibility could be a critical factor in developing next-generation BACE1 inhibitors with improved efficacy and reduced side effects. Further structural and functional studies are needed to elucidate these dynamic interactions and guide the optimization of these promising lead compounds for Alzheimer’s disease treatment.

**Fig 5:**
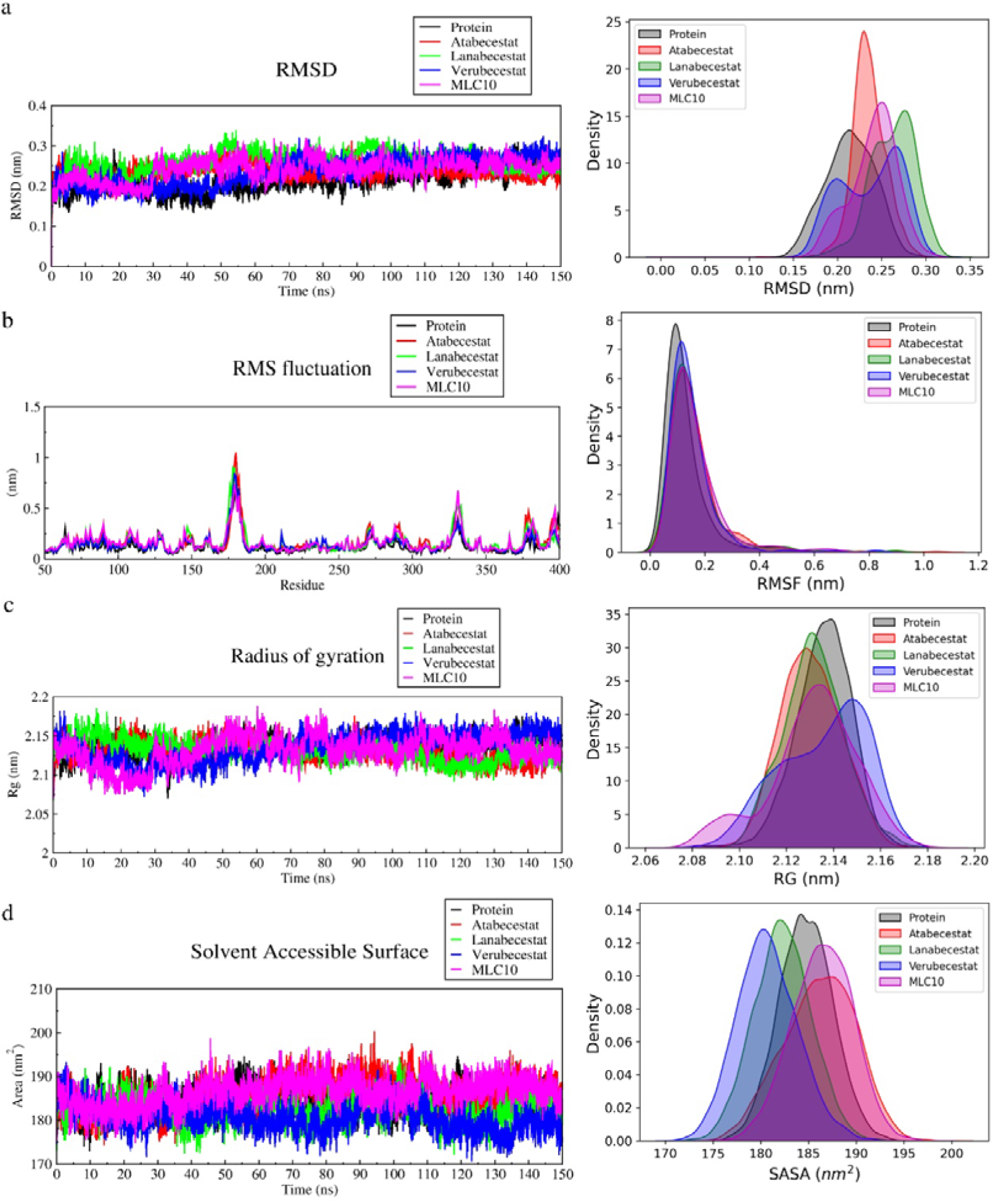
Molecular Dynamic Simulation analysis for 150 ns duration. (a) Root Mean Square Deviation (b) Root Mean Square Fluctuation (c) Radius of Gyration (d) Solvent Accessible Surface Area. Apo form of BACE1 is represented by black, in association with Atabecestat as red, in association with Lanabecestat as green, in association with Verubecestat as blue and in association with MLC10 as magenta

#### Root Mean Square Fluctuation (RMSF)

The RMSF analysis of BACE1 in complex with the respective ligands revealed intricate patterns of protein flexibility, shedding light on the dynamic behaviour of key functional regions and potential mechanisms of inhibition. The active site residues displayed relatively low RMSF values across all simulations, indicating a stable catalytic core. However, subtle differences emerged, with MLC10 inducing slightly higher fluctuations in this region compared to the established inhibitors (Fig 5b, magenta). This increased flexibility may enable dynamic interactions that optimize binding while maintaining the stability of the catalytic machinery. The flap region (residues 67-75) exhibited more pronounced variations in flexibility. In the protein-only simulation (Fig 5b, black), moderate fluctuations were observed, while Verubecestat appeared to stabilize the flap region (Fig 5b, blue), potentially contributing to its efficacy. Remarkably, MLC10 induced enhanced flexibility in the flap, with RMSF peaks reaching approximately 0.35 nm, which could facilitate a more dynamic “flap opening” mechanism, potentially improving inhibitor access to the active site. The 10s loop showed variable flexibility depending on the ligand, with Lanabecestat (Fig 5b, green) reducing fluctuations relative to the protein-only simulation, suggesting a stabilizing effect. In contrast, MLC10 increased flexibility in this region, possibly propagating structural changes to nearby catalytic residues and indirectly modulating enzyme activity. One striking feature of the RMSF plot was the pronounced peak in the Insert A region across all ligand-bound simulations, with Atabecestat inducing the highest flexibility (RMSF ∼1.1 nm) (Fig 5b, red). This loop, located near the active site, may play a key role in inhibition by altering the geometry of the substrate-binding cleft or influencing the dynamics of adjacent catalytic residues. MLC10 demonstrated a unique capacity to selectively enhance flexibility in key regions, such as the flap and 113s loop, while maintaining stability in others, including the active site and Insert A. This dynamic modulation contrasts with the more uniform stabilization observed with inhibitors like Verubecestat (Fig 5b, distribution plot), suggesting that MLC10 may fine-tune BACE1’s conformational landscape to achieve potent inhibition. This balance between stabilizing and mobilizing different functional regions may allow MLC10 to achieve more effective inhibition while potentially reducing off-target effects. The dynamic interplay between flexibility and stability observed in these regions provides valuable insights for the rational design of next-generation BACE1 inhibitors, offering new opportunities in the development of therapeutic agents for Alzheimer’s disease.

#### Radius of Gyration (Rg)

The radius of gyration (Rg) analysis reveals compelling insights into the conformational dynamics of BACE1 in its apo form and when bound to various inhibitors. These 150 ns molecular dynamics simulation captures subtle yet significant differences in protein compactness and flexibility. The apo protein exhibits the most variable Rg profile, fluctuating between 2.07 and 2.18 nm (Fig 5c, black), indicative of its inherent conformational plasticity. In contrast, all inhibitor-bound states demonstrate a general stabilization effect, albeit with distinct patterns. Atabecestat (Fig 5c, red) and Lanabecestat (Fig 5c, green) induce similar Rg profiles, maintaining the protein in a relatively compact state (Rg ∼2.10-2.15 nm) with reduced fluctuations compared to the apo form. Verubecestat (Fig 5c, blue) shows a unique behaviour, initially inducing a more compact structure (Rg ∼2.08-2.12 nm in the first 70 ns) before transitioning to a slightly more expanded state (Rg ∼2.14-2.17 nm) in the latter half of the simulation. Strikingly, MLC10 (Fig 5c, magenta) elicits the most dynamic Rg profile among all ligand-bound states. It induces rapid fluctuations between compact (Rg ∼2.08 nm) and expanded (Rg ∼2.18 nm) conformations throughout the simulation. This unique pattern suggests that MLC10 may promote a previously unobserved “breathing” motion in BACE1, potentially allowing the enzyme to sample a broader conformational landscape (Fig 5c, distribution plot). These findings highlight the complex interplay between ligand binding and protein dynamics. While traditional inhibitors tend to stabilize BACE1 in a more rigid conformation, MLC10’s ability to induce controlled flexibility may represent a significant shift in inhibitor design. This dynamic modulation could potentially overcome the limitations faced by previous BACE1 inhibitors in clinical trials, offering a new avenue for therapeutic intervention in Alzheimer’s disease.

#### Solvent Accessible Surface Area (SASA)

The solvent accessible surface area (SASA) analysis of BACE1 in its apo form and bound to various inhibitors reveals striking differences in protein-solvent interactions and conformational dynamics over a 150 ns molecular dynamics simulation. This comprehensive comparison provides crucial insights into their distinct mechanisms of action. The apo protein exhibits moderate SASA fluctuations (180-190 nm²) (Fig 5d, black), reflecting its native flexibility. In contrast, the inhibitor-bound states demonstrate markedly different SASA profiles, indicative of their unique effects on BACE1’s conformational landscape. Notably, MLC10 induces a dramatic increase in SASA (185-195 nm²) throughout the simulation (Fig 5d, magenta), suggesting a significant expansion of the protein’s surface exposure. This enhanced solvent accessibility implies that MLC10 may promote a more “open” conformation of BACE1, potentially altering its catalytic behaviour through increased solvent-enzyme interactions. Atabecestat shows a biphasic SASA pattern, initially maintaining a compact structure (175-185 nm²) before transitioning to a more exposed state (185-195 nm²) after 70 ns (Fig 5d, red). This transition hints at a time-dependent conformational change that could be crucial for its inhibitory mechanism. Lanabecestat (Fig 5d, green) and Verubecestat (Fig 5d, blue) consistently maintain lower SASA values (175-185 nm²), indicating they stabilize BACE1 in a more compact conformation. This reduced solvent exposure suggests these inhibitors may function by limiting the enzyme’s conformational freedom and substrate accessibility. The stark contrast between MLC10 and the clinically tested inhibitors in their effects on BACE1’s SASA (Fig 5d, distribution plot) provides a potential explanation for their divergent efficacies. MLC10’s ability to maintain an expanded protein conformation may allow for more dynamic inhibition, potentially overcoming the limitations of rigid, compact states induced by traditional inhibitors. These findings highlight the critical role of protein dynamics and solvent interactions in BACE1 inhibition. The unique conformational modulation induced by MLC10 represents a new avenue for inhibitor design, suggesting that promoting controlled flexibility, rather than rigid constraint, may be key to developing more effective BACE1 inhibitors for Alzheimer’s disease treatment.

#### MMPBSA

The binding free energies (ΔG) of BACE1 with respective inhibitors were evaluated using the MM-PBSA approach (Fig 6). Among the tested inhibitors, MLC10 exhibited the most favourable binding affinity with the lowest ΔG value of approximately −30 kcal/mol, outperforming the FDA-approved drugs Atabecestat and Lanabecestat, which showed binding free energies of around −25 kcal/mol and −15 kcal/mol, respectively. Interestingly, Verubecestat demonstrated the least favourable interaction with a binding energy of −9.6 kcal/mol. In contrast, the novel molecule MLC10 showed superior affinity, suggesting it could potentially serve as a highly potent BACE1 inhibitor. These results highlight the significant potential of MLC10 as a next-generation BACE1 inhibitor, surpassing the binding efficiency of the currently available drugs. The enhanced interaction observed with MLC10 may stem from its optimized structural features designed to fit the active site of BACE1 more effectively, underscoring the promise of rational drug design in developing more efficient therapeutic agents against Alzheimer’s disease. Further in vitro and in vivo studies would be essential to validate the therapeutic potential of MLC10 in clinical settings.

**Fig 6:**
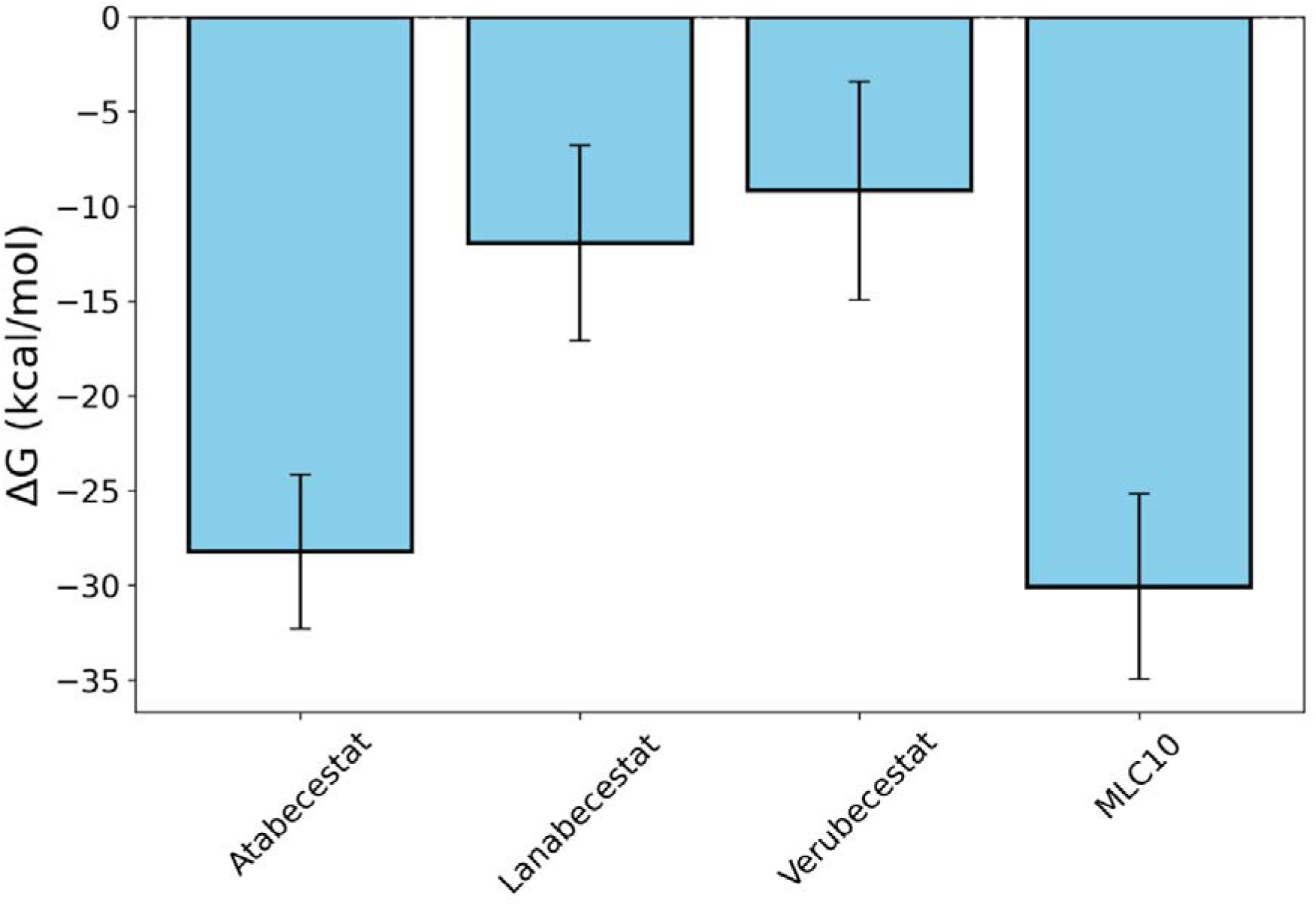
MMPBSA analysis of the known inhibitors and MLC10 for 150 ns MD Simulation. The error bar indicates the standard deviation

### Dynamic Cross Correlation Matrix (DCCM) Analysis

Dynamic Cross-Correlation Matrix (DCCM) analysis was conducted to evaluate the internal dynamics of BACE1 in its apo form and in complex with the inhibitors Atabecestat, Lanabecestat, Verubecestat, and the newly designed inhibitor MLC10 (Fig 7). The DCCM plots provide insights into residue-residue correlations, helping to elucidate the conformational flexibility and the dynamic coupling between critical regions, including the active site, flap region, and insert A, that are key for BACE1’s function. In its apo form, BACE1 exhibited moderate anti-correlated motions between the residues within the active site region (25-35, 225-235) (Fig 7a). Correlations between the active site and the flap region, which is known to regulate substrate access, were weakly anti-correlated, suggesting a semi-rigid, unengaged active site (Fig 7b, 7c). There was a lack of strong correlation between the active site and insert A (Fig 7d, 7e), highlighting the flexibility of the enzyme in the absence of an inhibitor. Binding of Atabecestat introduced stronger anti-correlated motions within the active site (Fig 7f), indicating some level of stabilization. However, the active site-flap dynamics (Fig 7g, 7h) remained similar to the apo state, with limited coupling between these regions. Active site 2 showed a stronger positive correlation with the flap region (Fig 7h) suggesting a coordinated movement of the domain in the presence of the inhibitor. The interaction with insert A also showed no significant enhancement in correlated motions (Fig 7i, 7j), suggesting that Atabecestat does not fully exploit the dynamic coupling between these functional regions. Lanabecestat binding resulted in a similar dynamic profile to Atabecestat, though with slightly enhanced moderate anti-correlation within the active site (Fig 7k). However, the active site-flap region coupling (Fig 7l, 7m) showed a similar profile of active site 1 – flap dynamics as that of Atabecestat but displayed a much stronger positive correlation in the active site 2 – flap dynamics (Fig 7m). The lack of significant interaction with insert A (Fig 7n, 7o) further corroborates the notion that Lanabecestat stabilizes BACE1 through direct binding without altering the enzyme’s global dynamic landscape. Verubecestat showed the weakest correlation within the active site (Fig 7p). The dynamic engagement of the flap region with active site 1 (Fig 7q) and active site 2 (Fig 7r) both displayed a dominance of positive correlation. The active site-insert A coupling (Fig 7s, 7t) was similarly weak, implying that Verubecestat binds with minimal perturbation to the enzyme’s overall dynamic behaviour. MLC10 displayed a markedly different dynamic profile compared to the other inhibitors. The dynamics of the active sites matched the dynamics shown by Lanabecestat with MLC10 showing slightly stronger positive correlation (Fig 7u), suggesting a tight, stabilizing interaction. Notably, MLC10 enhanced the coupling between the active site 1 and flap region (Fig 7V), with strong anti-correlations, implying that MLC10 effectively locks the flap in a closed conformation, potentially blocking substrate access more efficiently than the other inhibitors. Moreover, MLC10 displayed a similar dynamic of active site 2 with flap region as shown by Verubecestat, with slightly stronger positive correlation. This behaviour of MLC10 indicates that it replicates a similar dynamic of Verubecestat, that failed Clinical trials phase 3 [17]) with some nuanced differences that might prove MLC10 as a better inhibitor. Furthermore, MLC10 induced significant anti-correlated motions between the active site 1 and insert A (Fig 7x) that matches closely with the apo form dynamics (Fig 7d), a feature absent in the other complexes. Furthermore, it showed a strong anti-correlation between active site 2 and insert A with weak positive correlation between residues 225228 and 158-160 (Fig 7y). This suggests that MLC10’s binding extends beyond simple inhibition at the active site and influences the broader conformational dynamics of BACE1. By enhancing both the local and global dynamic coupling, MLC10 may offer superior inhibition by stabilizing the enzyme in a catalytically inactive conformation. By tightening the coupling between the active site and distal regions such as the flap and insert A, MLC10 may not only block substrate entry but also restrict the conformational flexibility necessary for BACE1’s catalytic function. These dynamic insights strongly support the potential of MLC10 as a next-generation therapeutic agent targeting BACE1 in Alzheimer’s disease treatment. Further studies, including experimental validation and structural analysis, will be critical to confirm this proposed mechanism.

**Fig 7:**
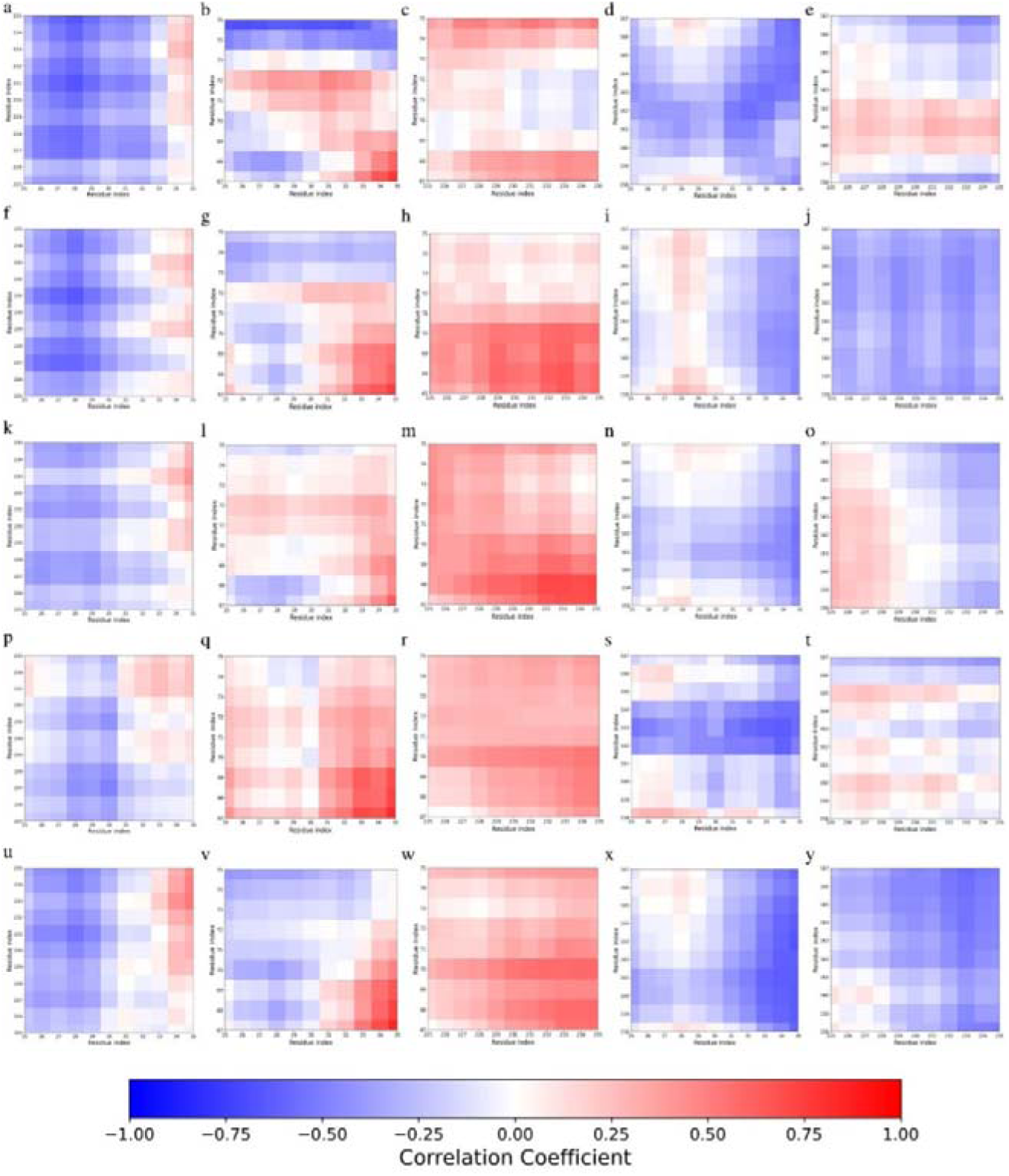
DCCM analysis for the 150 ns MD simulation of Apo form of BACE1 with known inhibitors and MLC10. BACE1 Apo (first row), in association with Atabecestat (second row), in association with Lanabecestat (third row), in association with Verubecestat (forth row), in association with MLC10 (fifth row). The first column corresponds to DCCM between active site 1 (includes Asp32) and active site 2 (includes Asp228), the second column corresponds to DCCM between active site 1 and flap region (residues 65-75), the third column corresponds to DCCM between active site 2 and flap region, the fourth column corresponds to DCCM between active site 1 and Insert A (residues 158-167) and the second column corresponds to DCCM between active site 2 and Insert A. The colour bar denotes the correlation coefficient with blue corresponding to anti-correlation and red corresponding to positive correlation

### Principal Component Analysis

The Free Energy Landscapes and associated structural conformations of BACE1 in complex with Atabecestat, Lanabecestat, Verubecestat, and MLC10 reveal distinct ligand-induced conformational behaviours, offering profound insights into the mechanism of BACE1 inhibition. Atabecestat (Fig 8a) induces a FEL with two well-defined energy minima, suggesting a bimodal conformational distribution. The structural overlay shows differences in Insert F and Insert D regions between the two stable states (Fig 8a, overlapped structure). This conformational plasticity may contribute to Atabecestat’s mechanism of action, potentially allowing it to modulate BACE1 function through dynamic conformational selection. Lanabecestat (Fig 8b) generates a more complex energy landscape with a diffused local minimum, indicating that it allows BACE1 to sample a diverse range of conformations. The structural comparison between the two most stable states (Fig 8b, overlapped structure) reveals subtle yet significant differences, particularly in the positioning of flap regions and slight shifts in Insert D. This conformational heterogeneity could be linked to Lanabecestat’s efficacy profile and potential side effects. Verubecestat (Fig 8c) exhibits three distinct local minima, suggesting it allows more conformational flexibility in BACE1. The structural overlay (Fig 8c, overlapped structure) shows less dramatic differences between the two stable states compared to Atabecestat and Lanabecestat, with the main variations observed in Insert F and a minor difference in Insert D region. This increased flexibility might affect Verubecestat’s inhibitory mechanism and could potentially explain its clinical profile. Intriguingly, MLC10 (Fig 8d) produces a similar energy landscape, like Verubecestat, with a single, deep global minimum and two few smaller local minima. The structural overlay (Fig 8d, overlapped structure) of the two most stable conformations shows remarkable similarity, with only minor differences in Insert A region. This suggests that MLC10 significantly stabilizes a particular conformation of BACE1, potentially leading to more potent and specific inhibition. The structural similarity between its stable states indicates a more consistent binding mode, potentially leading to more predictable and reliable inhibition. These structural changes suggest that MLC10 may achieve superior inhibition by locking BACE1 into a specific conformation that is incompatible with its catalytic function. This conformational restriction could potentially lead to improved efficacy and reduced side effects compared to the other ligands.

**Fig 8:**
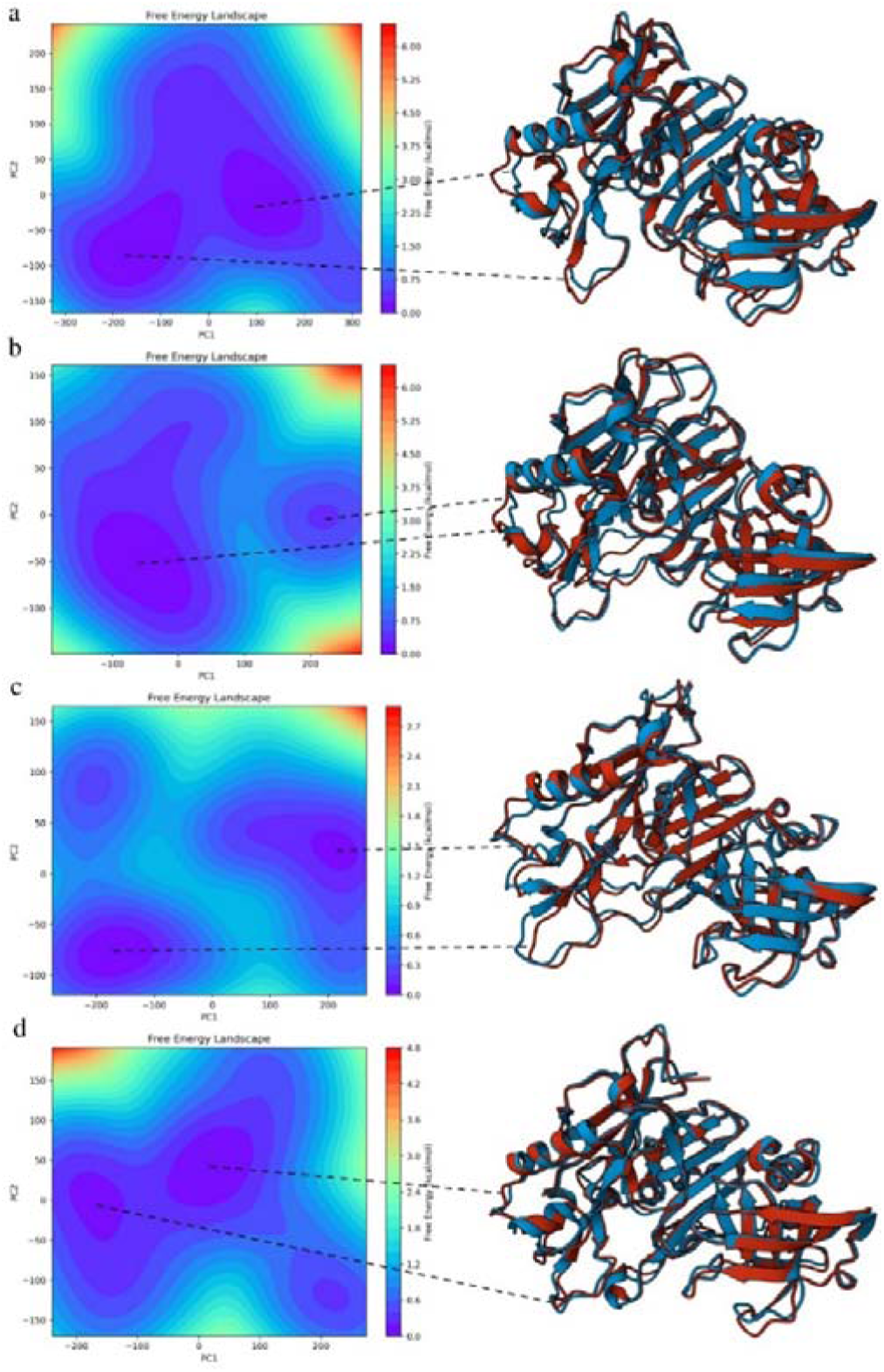
PCA based Free Energy Landscape (FEL) Analysis of BACE1 with known inhibitors and MLC10. (a) Atabecestat (b) Lanabecestat (c) Verubecestat (d) MLC10. The right panel shows the overlap of representative structure of the energetically minimised basin with red coloured structure corresponding to most energetically stable and blue represents the second energetically minimised structure

### Deformity Analysis

Deformity analysis of BACE1 in complex with various inhibitors provides crucial insights into the molecular mechanisms of enzyme inhibition and ligand specificity. The results reveal distinct patterns of protein deformation induced by each ligand, offering a nuanced understanding of BACE1 inhibition that goes beyond simple active site occupancy. Atabecestat (Fig 9a) induces significant localized deformations in BACE1, particularly evident in residues 70-80 (some portion of flap region), 300-310 (prior to Insert F), and 330-340. The overall deformation pattern closely mirrors that of the apo protein, suggesting that Atabecestat allows BACE1 to maintain much of its native flexibility. Lanabecestat (Fig 9b) shows a similar overall deformation profile to Atabecestat, but with notable differences. It induces higher deformity in residues 70-80, 190-210, 230-240 and 330-340 region while slightly reducing flexibility in the 300-320 range. This unique pattern may explain Lanabecestat’s distinct pharmacological profile. Verubecestat (Fig 9c) presents a markedly different deformation landscape. It significantly reduces deformity in residues 70-80, 300-320 while enhancing flexibility in other regions including residues 110-120, 155-175, 190-200. This redistribution of protein flexibility could be key to Verubecestat’s mechanism of action, potentially locking flap region (residues 70-80) and Insert F (residues 311-317) while allowing compensatory movements of 113S loop (residues 110-117) and Insert A (residues 158-167). MLC10 (Fig 9d) exhibits the most intriguing deformation profile. It induces pronounced, localized deformations that are distinct from the apo protein’s natural flexibility. Notably, MLC10 causes sharp spikes in deformity in flap region (residues 70-80), 113S loop (residues 110-117), and Insert A (residues 158-167), while dampening flexibility in Insert F (residues 311-317) and Insert D (residues 270-274). This pattern suggests a highly specific interaction mode that reshapes BACE1’s energy landscape in a unique manner. The distinct deformation pattern, particularly in regions away from the presumed active site, hints at potential allosteric mechanisms. MLC10 may be leveraging long-range conformational changes to achieve inhibition, a sophisticated mechanism that could lead to improved efficacy. Moreover, the sharp deformation peaks suggest that MLC10 might be selecting and stabilizing specific BACE1 conformations that are particularly unfavourable for catalytic activity. Unlike the other inhibitors, which largely work within the protein’s native flexibility landscape, MLC10 appears to reshape this landscape more dramatically. This could result in a more effective “conformational freeze” of BACE1, potentially leading to superior inhibition.

**Fig 9:**
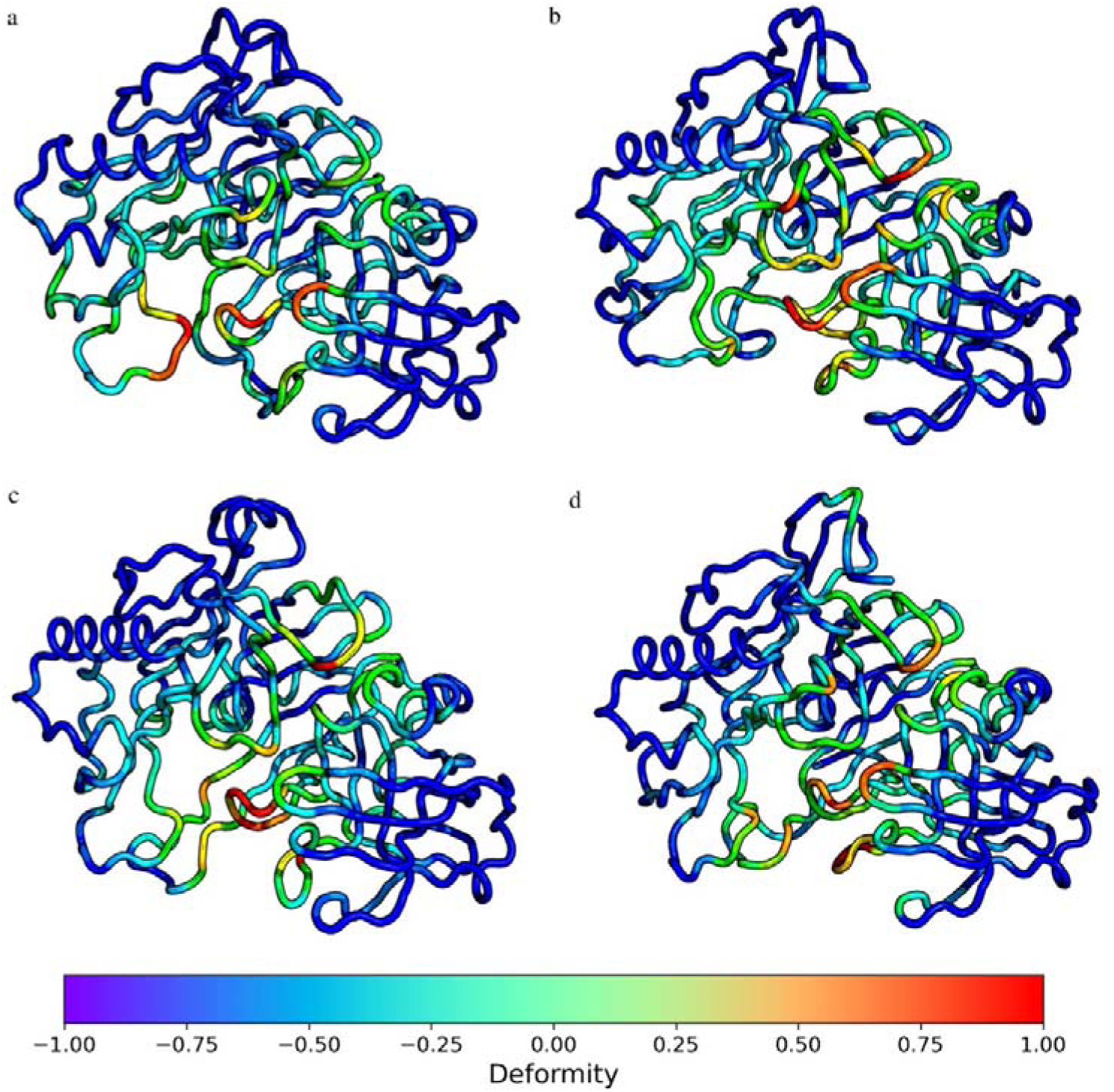
Deformity analysis of BACE1 with known inhibitors and MLC10. (a) Atabecestat (b) Lanabecestat (c) Verubecestat (d) MLC10. The colour bar corresponds to the extent of deformity with blue representing structured region and red corresponding to highly deformed region

### Normal Mode Energy Fluctuation Analysis

The normal mode analysis (NMA) of BACE1 in complex with Atabecestat, Lanabecestat, Verubecestat, and MLC10 provides an in-depth understanding of the global dynamic behaviour of the protein and its energy landscape under the influence of each inhibitor. By examining energy fluctuations, which reflect the collective movements of residues, we can infer how each inhibitor influences the intrinsic flexibility of BACE1 and its potential to stabilize or destabilize key functional regions, such as the active site, flap, and insert A. The energy fluctuation profile for the Atabecestat-BACE1 (Fig 10a) complex reveals significant peaks in energy fluctuations at residues 270-275 (Insert D region) and 310-320 (Insert F), similar to the RMSF fluctuations. These peaks indicate that Atabecestat fails to adequately stabilize the dynamic regions of the protein. The elevated energy fluctuations suggest that the inhibitor does not effectively suppress the inherent mobility of BACE1, particularly in areas crucial for catalytic activity. The active site residues (Asp32 and Asp228) exhibit moderate energy fluctuations, suggesting that Atabecestat provides only partial stabilization of the catalytic region. The Lanabecestat-BACE1 (Fig 10b) complex shows a slight reduction in energy fluctuations compared to Atabecestat, with moderate peaks still present in the flap region. Moreover, Lanabecestat reduces the fluctuation in Insert A (residues 158-167) and Insert F (residues 311-317) as compared to the apo form and Atabecestat. While Lanabecestat provides a marginally better stabilization of the active site and surrounding regions, the energy fluctuations suggest that this inhibitor also leaves key dynamic regions with substantial mobility. The failure to fully stabilize the flap region may allow for substrate access, thereby reducing its inhibitory efficacy. The energy fluctuations in the active site remain moderate, implying that Lanabecestat, like Atabecestat, only partially restrains BACE1’s conformational flexibility. Verubecestat demonstrates (Fig 10c) a more favourable energy fluctuation profile, with slightly higher peak in the flap and Insert D region compared to Lanabecestat. However, the fluctuations in Insert A region remain high, indicating residual flexibility in this distal region. While Verubecestat offers better stabilization of the flap, the persistent energy fluctuations in Insert A suggest that it does not fully lock down the conformational states required for BACE1 inhibition. This partial stabilization could indicate its failure and limitations in phase 3 clinical applications. The active site shows moderate-to-low energy fluctuations, indicating improved but incomplete stabilization compared to the other inhibitors. MLC10 (Fig 10d) exhibits a unique fluctuation profile, especially in key functional regions such as the flap and Insert A. The reduction in energy fluctuations in the flap region as compared to Verubecestat suggests that MLC10 effectively stabilizes this critical area, locking it into a closed conformation that hinders substrate access. Furthermore, the slight increment of energy fluctuations in Insert A indicates that MLC10 follows a very close fluctuation profile as that of Verubecestat, yet induces distinct dynamics. The active site residues, particularly Asp32 and Asp228, display minimal energy fluctuations, indicating that MLC10 provides strong stabilization of the catalytic core. Moreover, MLC10 induces a higher fluctuation in 10S loop region (residues 5-16), Insert D (residues 270-274) and distal region (residues 360-370). By suppressing the inherent flexibility of local region while increasing the distal region fluctuation, MLC10 likely inhibits BACE1’s ability to undergo the conformational changes necessary for substrate binding and catalysis. The ability of MLC10 to reduce energy fluctuations in the active site and distal regions, such as the flap and insert A (similar to Verubecestat), suggests that it induces a more rigid and stabilized conformation of BACE1. This rigidity is likely responsible for its enhanced inhibitory effect, as it prevents the dynamic transitions required for BACE1’s catalytic function. The combination of strong local and global stabilization of BACE1 by MLC10 sets it apart from the other inhibitors. This enhanced stabilization, particularly in regions that are critical for enzymatic activity, positions MLC10 as a promising candidate for further development in the treatment of Alzheimer’s disease.

**Fig 10:**
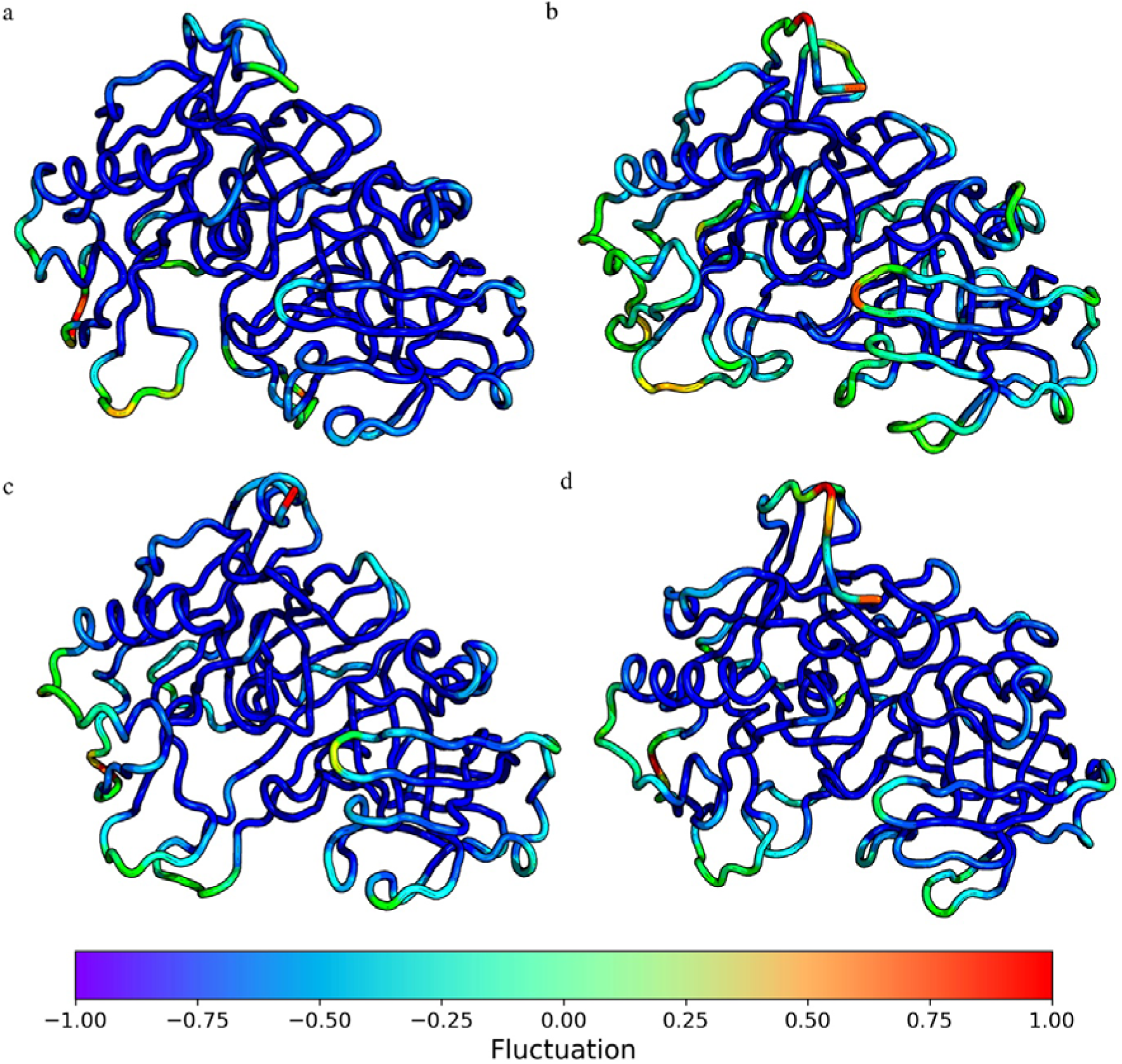
NMA based Fluctuation analysis of BACE1 with known inhibitors and MLC10. (a) Atabecestat (b) Lanabecestat (c) Verubecestat (d) MLC10. The colour bar corresponds to the extent of fluctuation with blue representing energetically least fluctuating region and red corresponding to energetically highly fluctuating region

## Discussion

Alzheimer’s disease (AD) represents one of the most pressing challenges in modern medicine, characterized by progressive cognitive decline and significant socioeconomic burdens worldwide [1,2]. At the molecular level, the pathogenesis of AD is intricately linked to the accumulation of neurotoxic amyloid-beta (Aβ) peptides, which are generated through the cleavage of amyloid precursor protein (APP) by β-site amyloid precursor protein cleaving enzyme 1 (BACE1) [43,44,45,46]. As a pivotal enzyme in the amyloidogenic pathway, BACE1 has emerged as a prime target for therapeutic intervention aimed at reducing Aβ production and mitigating the progression of AD. The current study presents a significant advancement in the quest for effective BACE1 inhibitors, introducing MLC10 as a top candidate that exhibits a unique mechanism of action. Unlike traditional inhibitors that stabilize BACE1 in rigid conformations, MLC10 promotes controlled flexibility within the enzyme. This dynamic modulation is crucial, as it allows for a more nuanced interaction with the substrate, potentially enhancing the inhibitor’s efficacy while minimizing off-target effects. The findings reveal that MLC10 induces specific conformational changes in BACE1, as evidenced by the substantial increase in solvent accessible surface area (SASA) observed during molecular dynamics simulations. This suggests that MLC10 may facilitate a more “open” conformation of BACE1, thereby altering its catalytic behaviour and enhancing solvent-enzyme interactions. Moreover, the study’s exploration of MLC10’s effects on various regions of BACE1, particularly those distant from the active site, hints at potential allosteric mechanisms of inhibition. This is a noteworthy departure from conventional inhibitor design, which typically focuses on direct active site interactions. By reshaping the energy landscape of BACE1, MLC10 may stabilize conformations that are less favourable for catalytic activity, thereby achieving a more effective “conformational freeze” of the enzyme.

The integration of machine learning into the drug discovery pipeline further enhances the significance of this work. By leveraging advanced algorithms to navigate the vast chemical space and predict therapeutic outcomes, the study exemplifies how modern computational techniques can accelerate the identification of promising candidates. This approach not only streamlines the drug development process but also increases the likelihood of success in clinical trials, which have historically faced high failure rates in AD research.

In conclusion, the current work represents a pivotal step forward in the development of BACE1 inhibitors for Alzheimer’s disease. By elucidating the unique mechanism of action of MLC10 and emphasizing the importance of protein dynamics, this study lays the groundwork for future therapeutic strategies that prioritize flexibility and allosteric modulation. As the field of Alzheimer’s research continues to evolve, the insights gained from this study will be invaluable in guiding the rational design of more effective and targeted interventions, ultimately contributing to the fight against this devastating disease.

