## Supplementary file for "Machine Learning-Enhanced Drug Discovery for BACE1: A Novel Approach to Alzheimer’s Therapeutics"

Satyam Sangeet^1*^

^1^CompObelisk, Makolia, Bahraich, Uttar Pradesh, India – 271802

**The current supplementary files include:**

- Table displaying the structure of the generated ligands using DrugGPT
- Table displaying the Lipinski’s RO5 for the generated ligands
- Figure displaying the distribution analysis of the Lipinski’s RO5 for the generated ligands
- Figure displaying the Tanimoto Similarity for the known inhibitors and the generated ligands
- Figure displaying redocked structures of 8AP overlapped with each other
- Figure displaying the triplicate analysis of apo form of BACE1 for 150 ns MD Simulation
- Figure displaying the triplicate analysis of BACE1 with Atabecestat for 150 ns MD Simulation
- Figure displaying the triplicate analysis of BACE1 with Lanabecestat for 150 ns MD Simulation
- Figure displaying the triplicate analysis of BACE1 with Verubecestat for 150 ns MD Simulation
- Figure displaying the triplicate analysis of BACE1 with MLC10 for 150 ns MD Simulation
- Figure displaying the MD Analysis statistics for the inhibitors
- Figure displaying the DCCM analysis for the apo form of BACE1 and in association with respective inhibitors
- Figure displaying the DCCM analysis of active site 1 and active site 2 with 10s loop and 113s loop for the apo form of BACE1 and in association respective inhibitors
- Figure displaying the Normal mode variance for apo form of BACE1 and in association with respective inhibitors
- Figure displaying the deformity analysis of apo form of BACE1 in comparison with the respective inhibitors
- Figure displaying the NMA based fluctuation analysis of apo form of BACE1 in comparison with the respective inhibitors

Table S1: Chemical structure of the compounds generated by DrugGPT against BACE1

| Compound | | | Structure |
| --- | --- | --- | --- |
| C1 | | 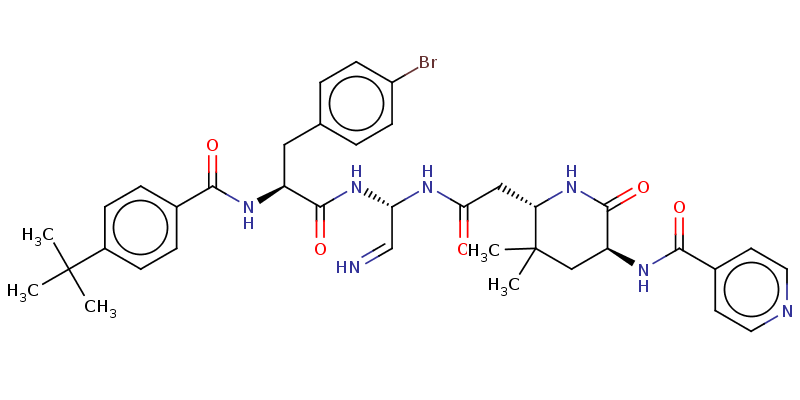 | |
| C2 | 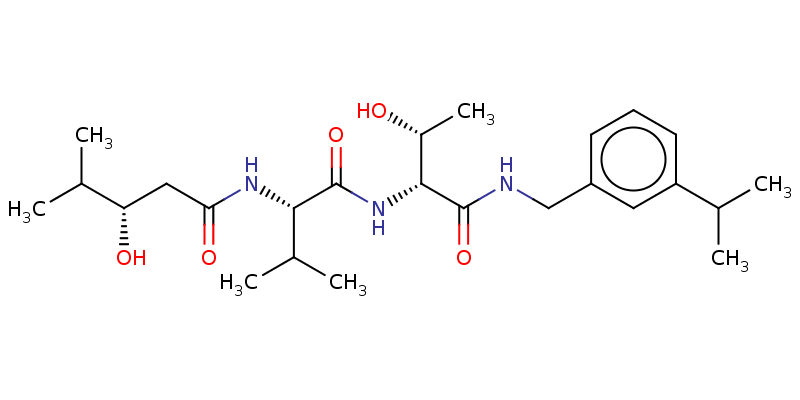 | | |
| C3 | 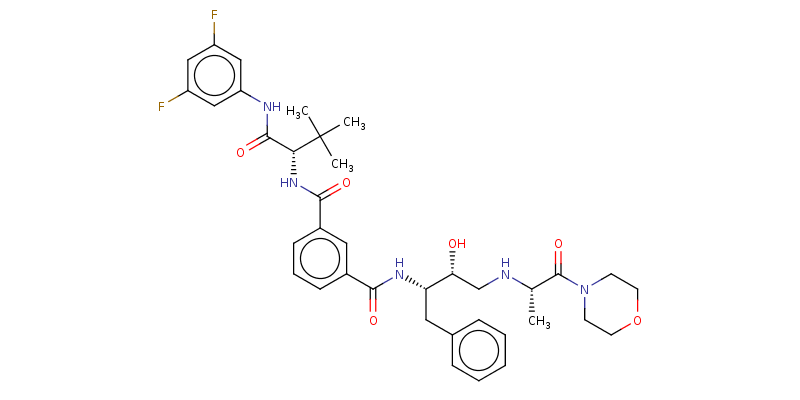 | | |
| C4 | 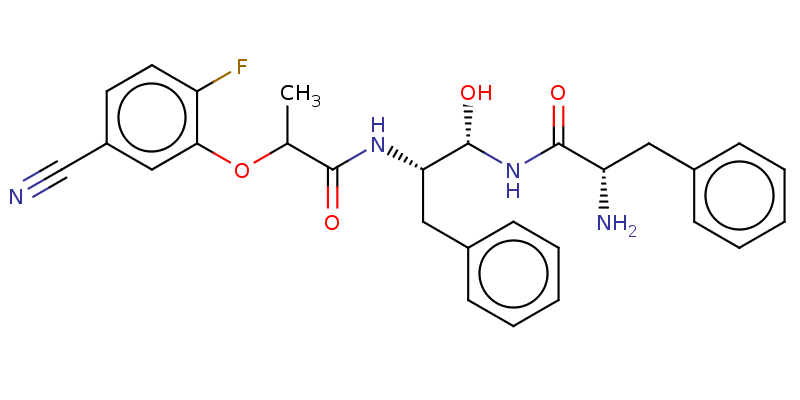 | | |
| C5 | 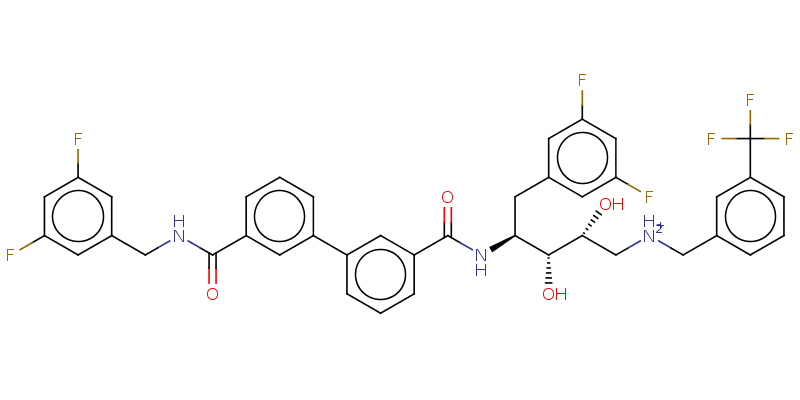 | | |
| C6 | 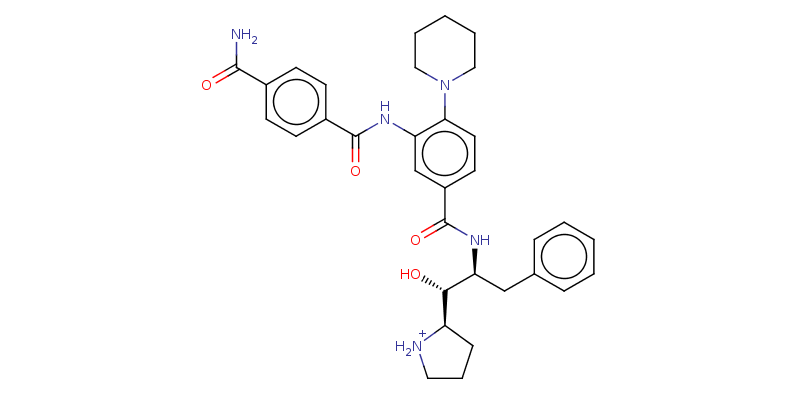 | | |
| C7 | 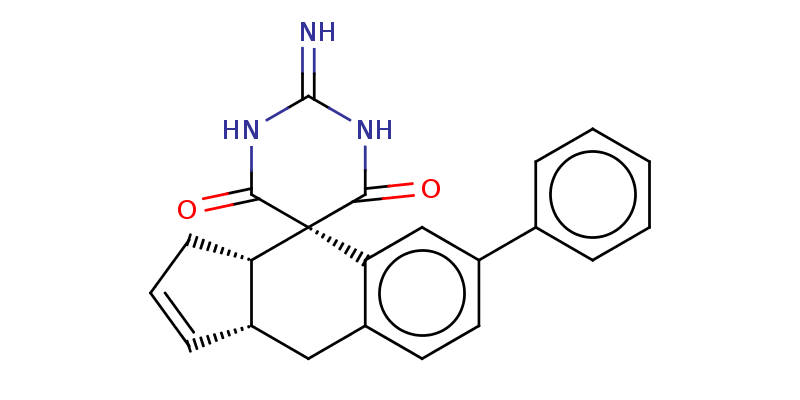 | | |
| C8 | 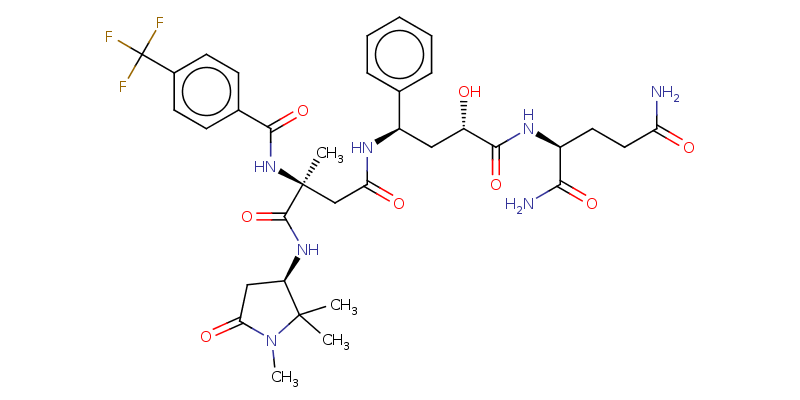 | | |
| C9 | 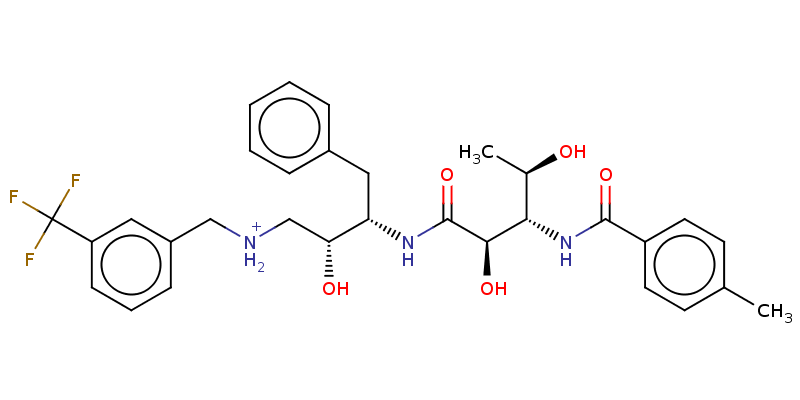 | | |
| C12 | 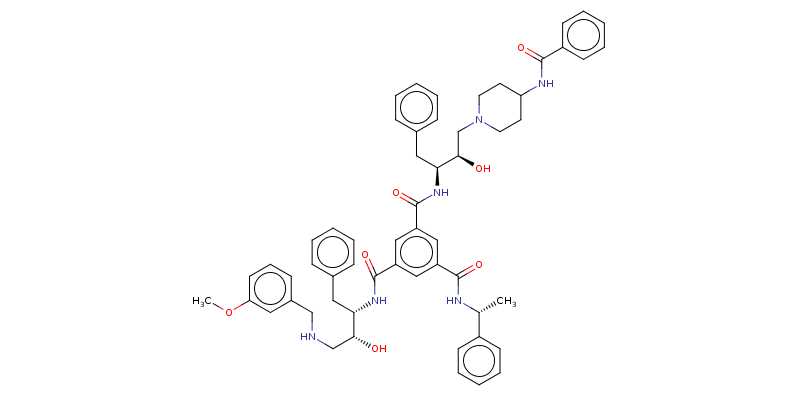 | | |
| C13 | 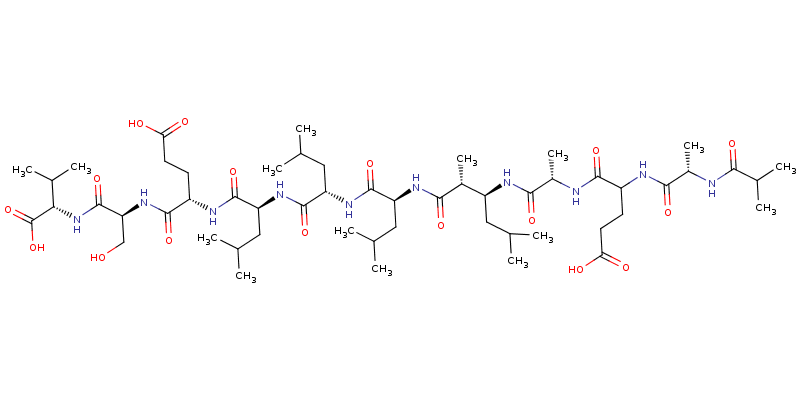 | | |
| C14 | 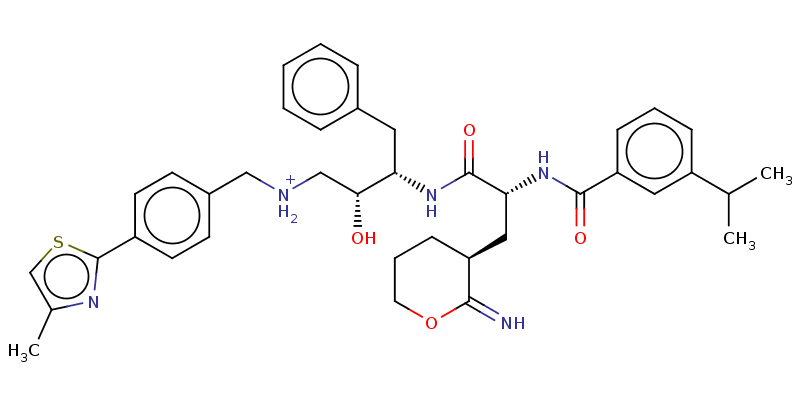 | | |
| C15 | 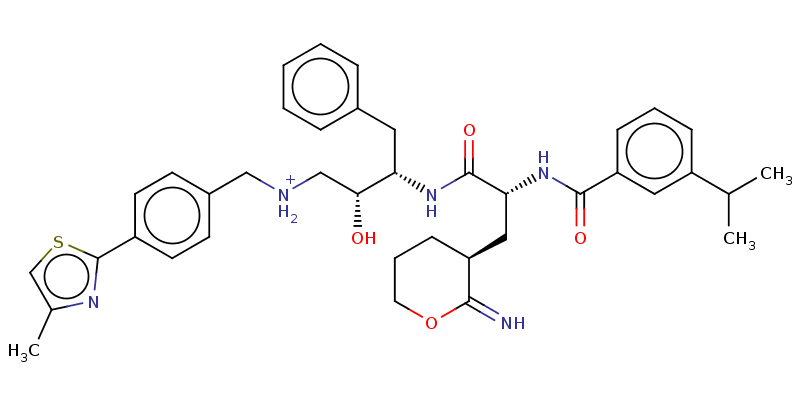 | | |
| C16 | 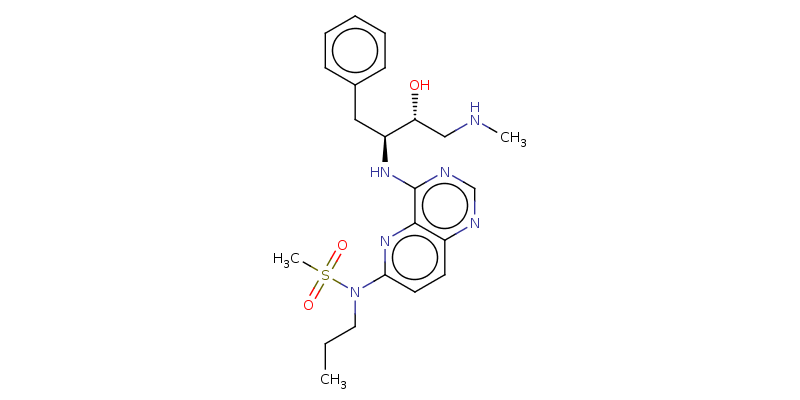 | | |
| C17 | 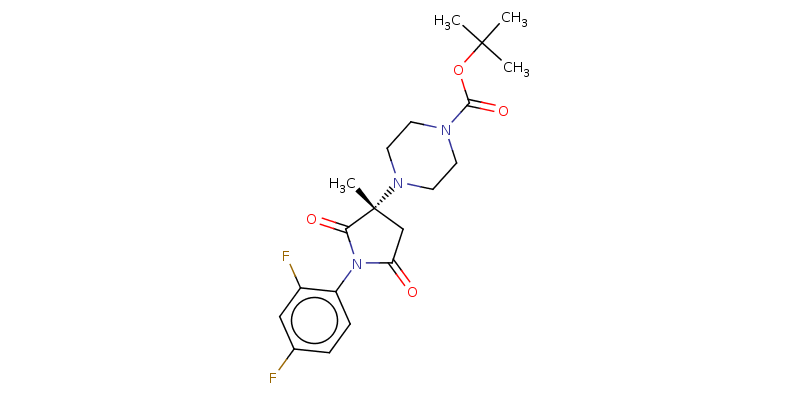 | | |
| C18 | 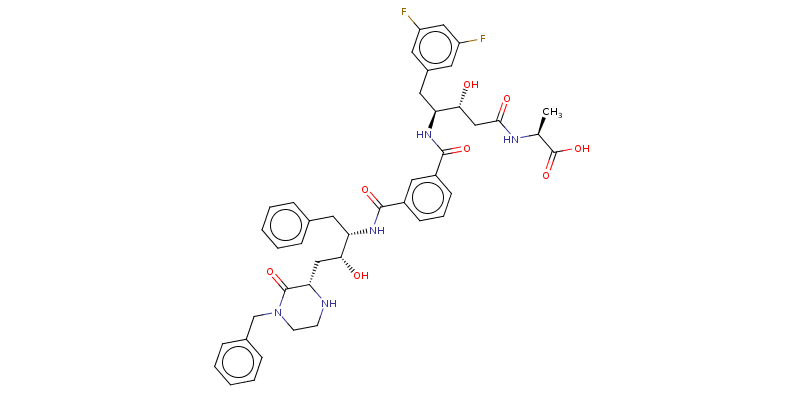 | | |
| C19 | 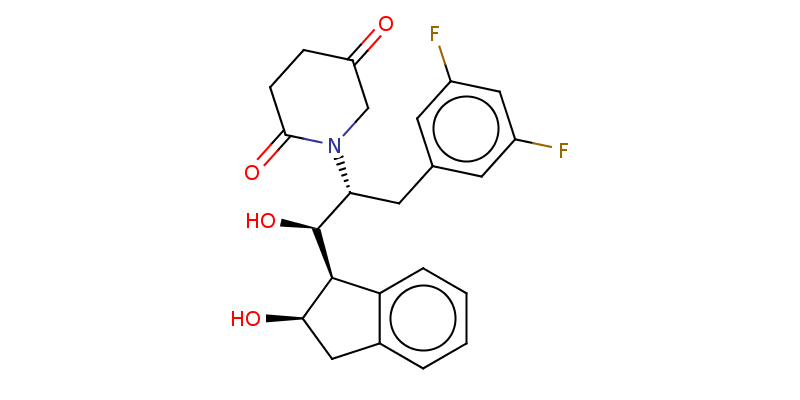 | | |
| C20 | 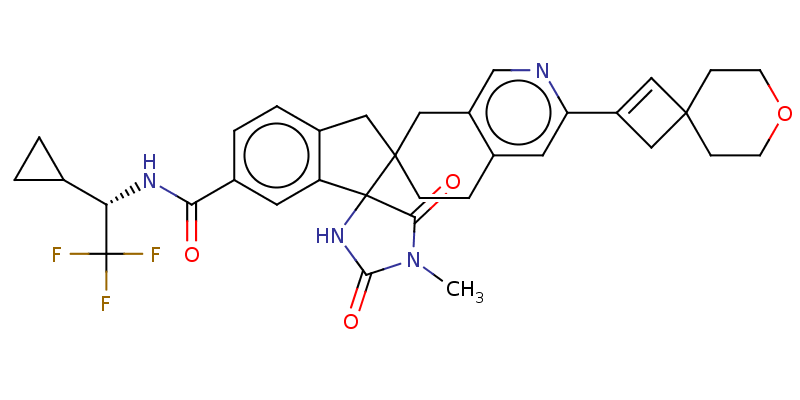 | | |
| C21 | 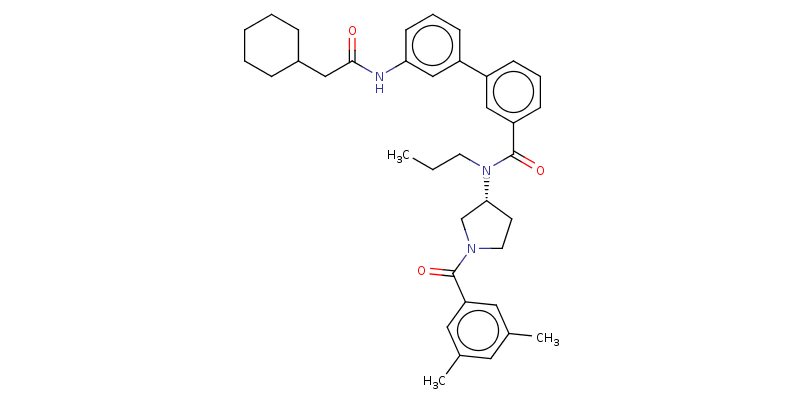 | | |
| C22 | 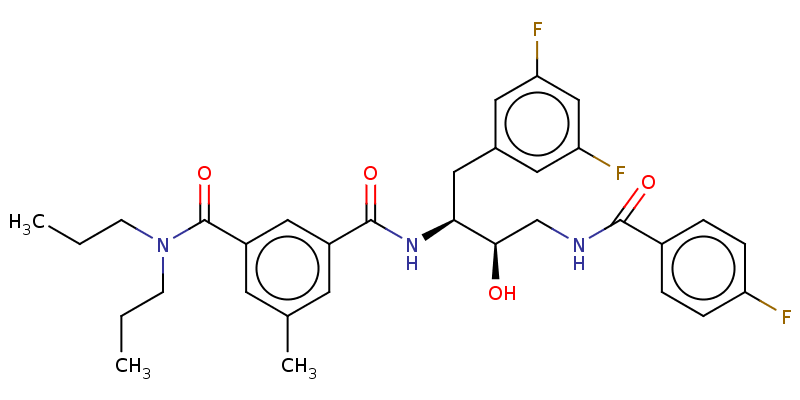 | | |
| C23 | 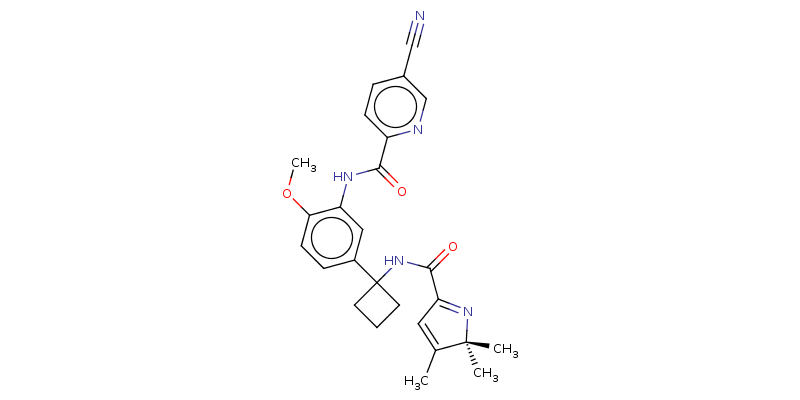 | | |
| C24 | 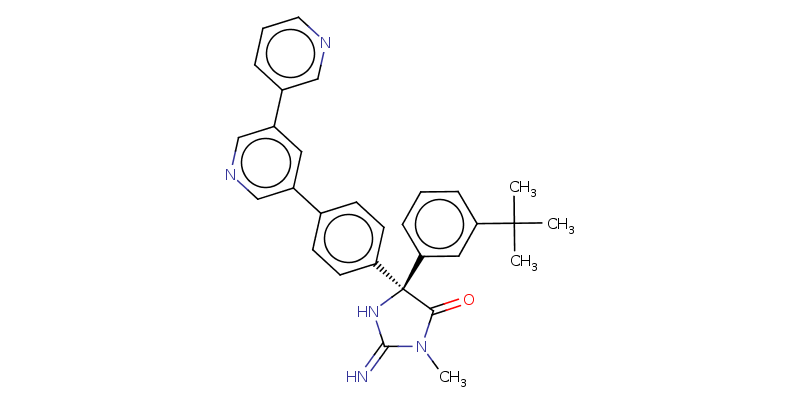 | | |
| C25 | 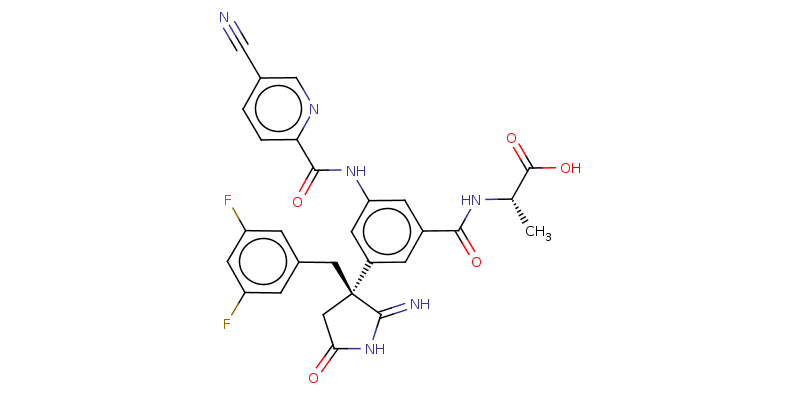 | | |
| C26 | 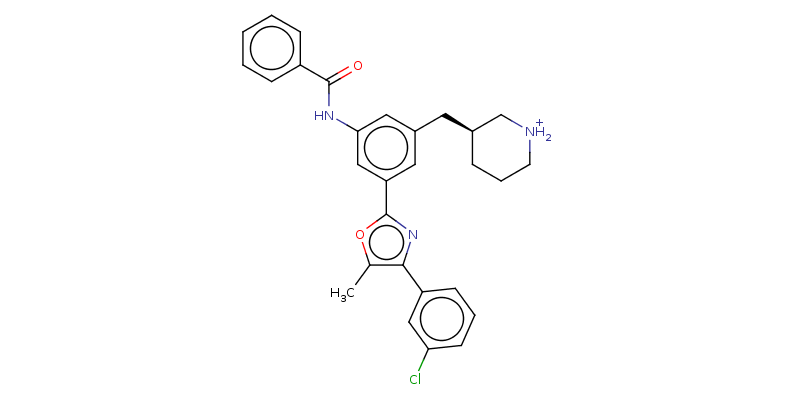 | | |
| C27 | 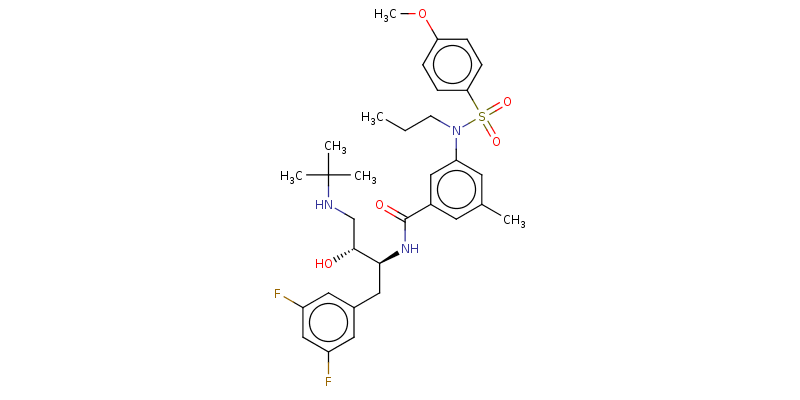 | | |
| C28 | 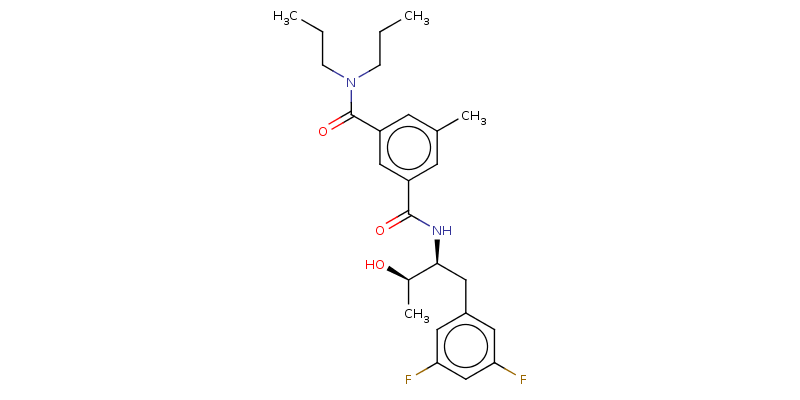 | | |
| C29 | 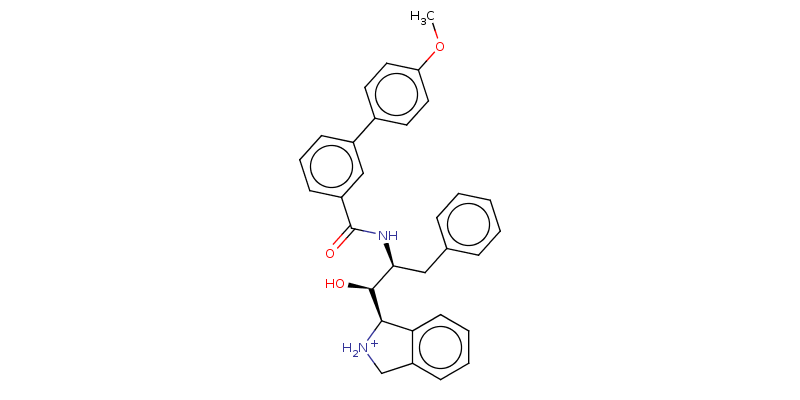 | | |
| C30 | 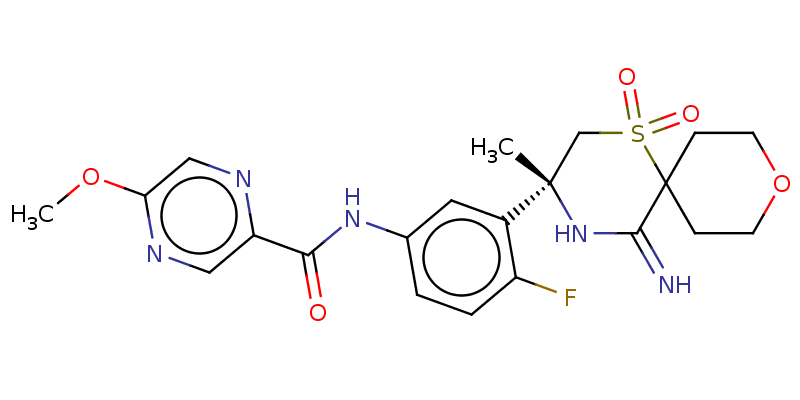 | | |
| C31 | 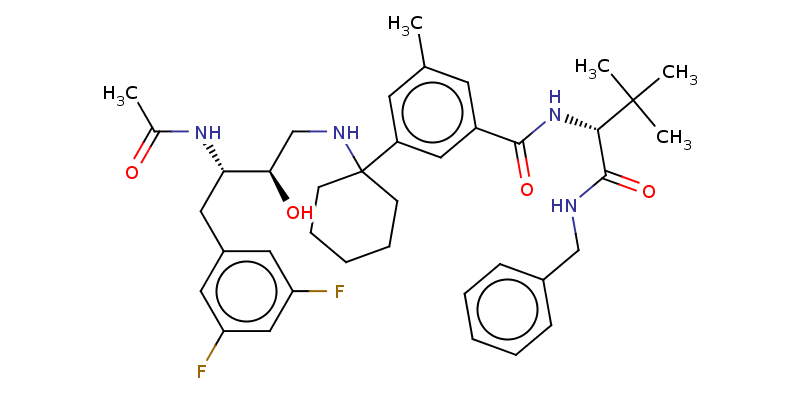 | | |
| C32 | 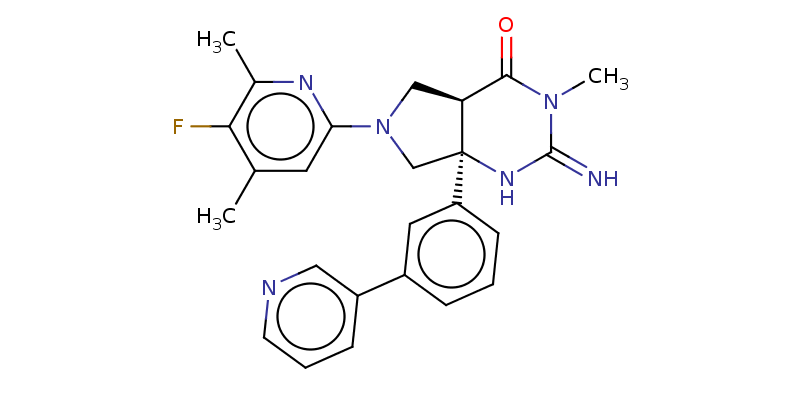 | | |
| C33 | 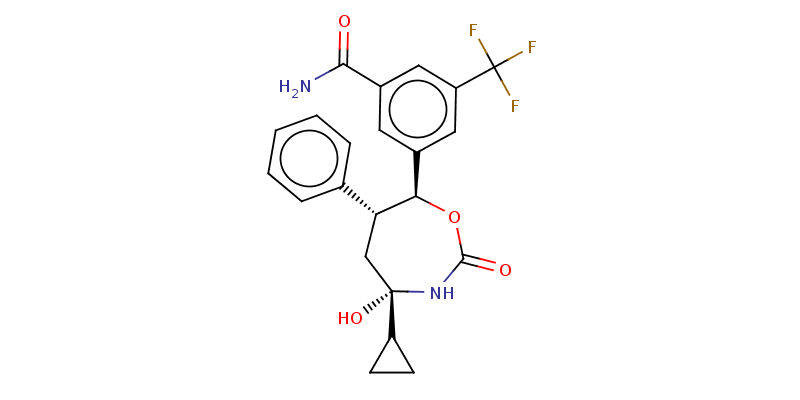 | | |
| C34 | 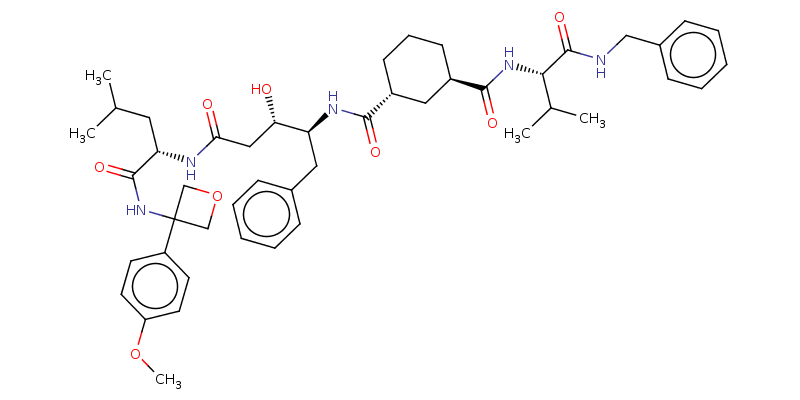 | | |
| C35 | 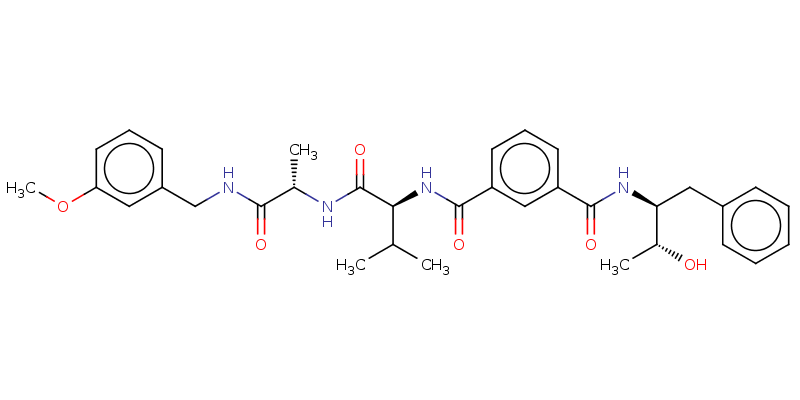 | | |
| C36 | 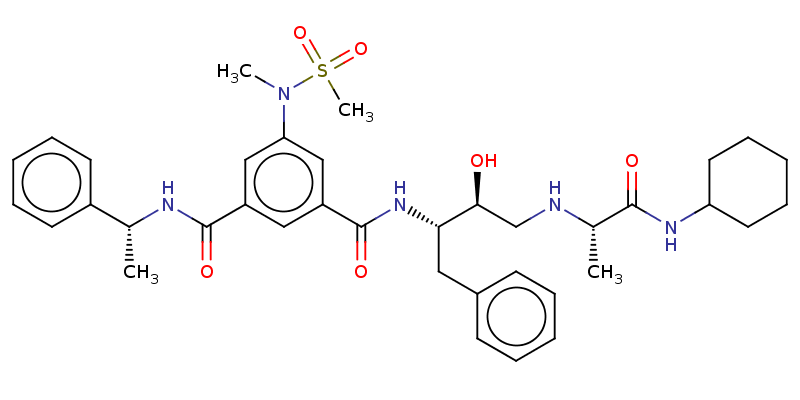 | | |
| C37 | 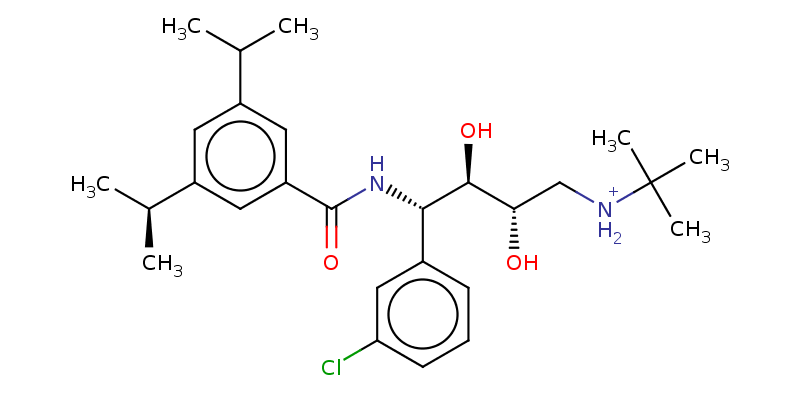 | | |
| C38 | 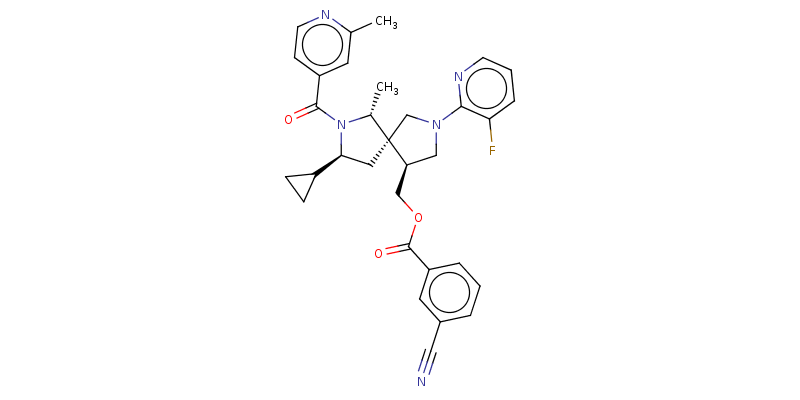 | | |
| C39 | 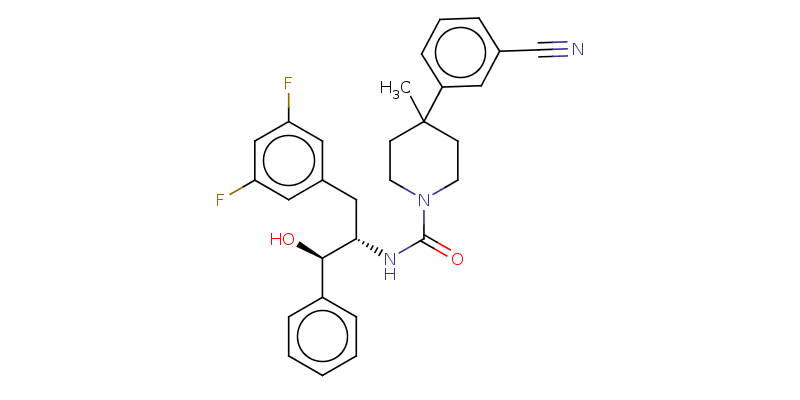 | | |
| C40 | 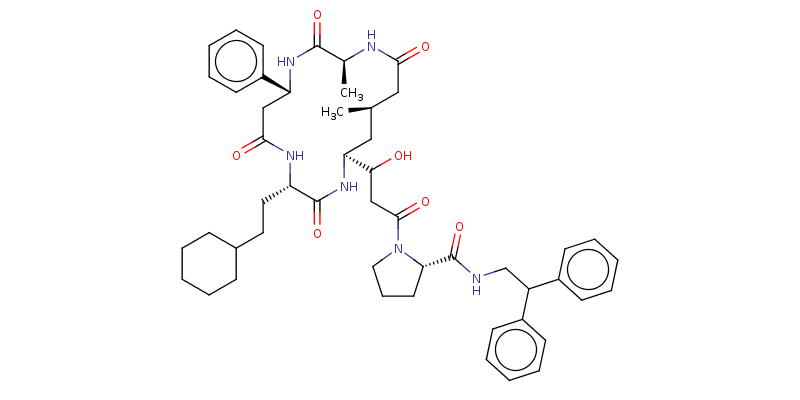 | | |
| C41 | 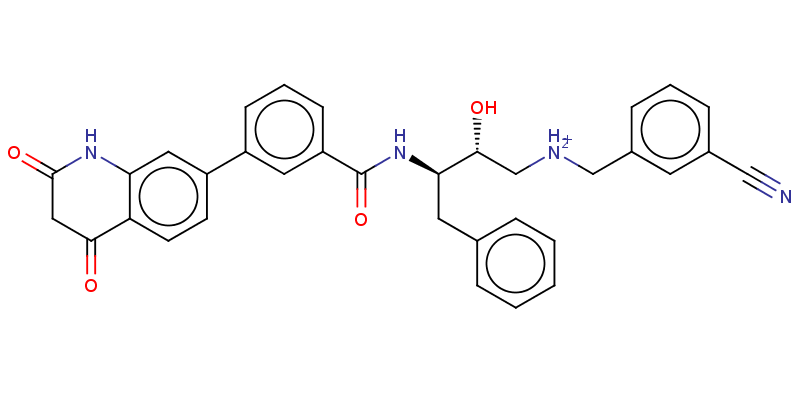 | | |
| C42 | 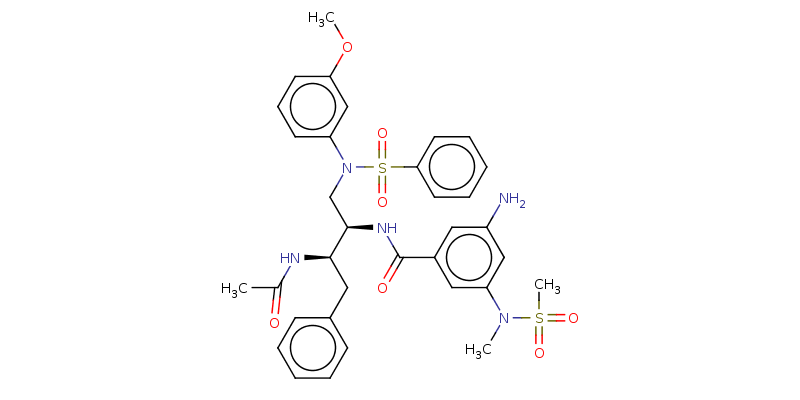 | | |
| C43 | 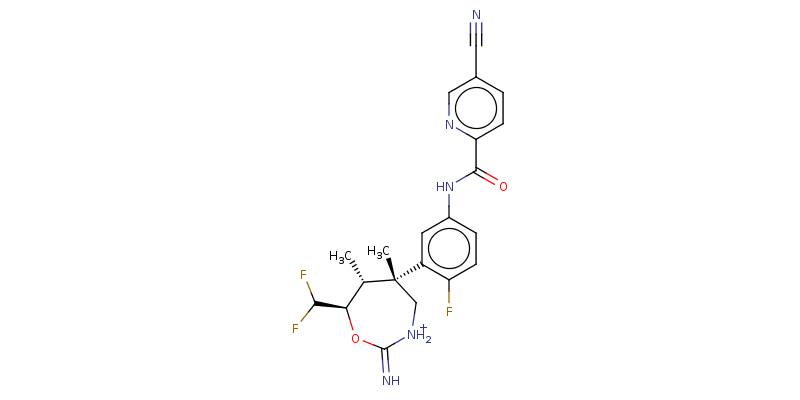 | | |
| C44 | 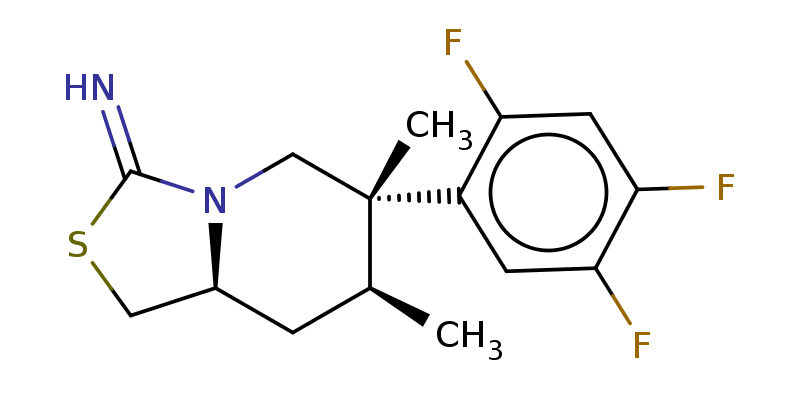 | | |
| C45 | 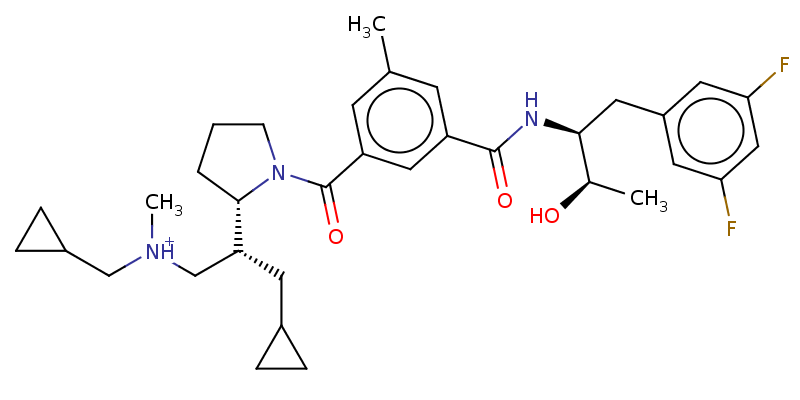 | | |
| C46 | 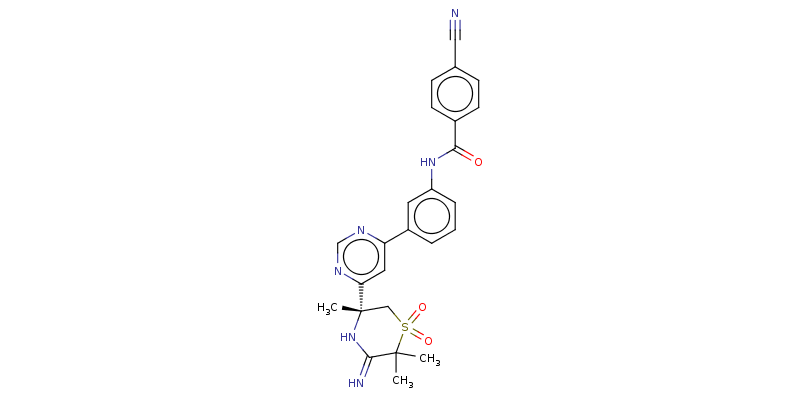 | | |
| C47 | 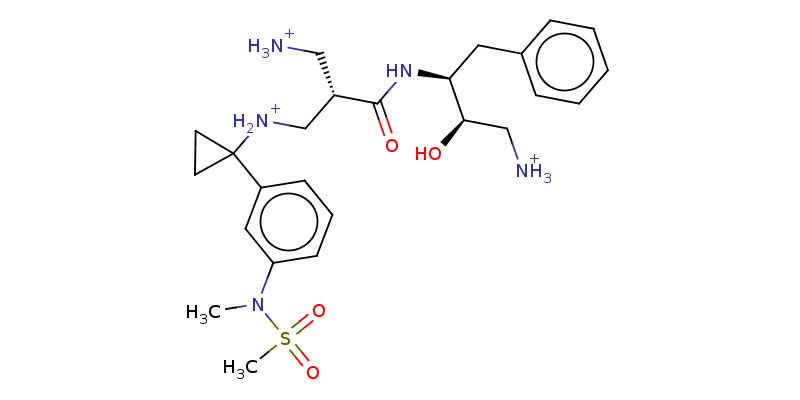 | | |
| C48 | 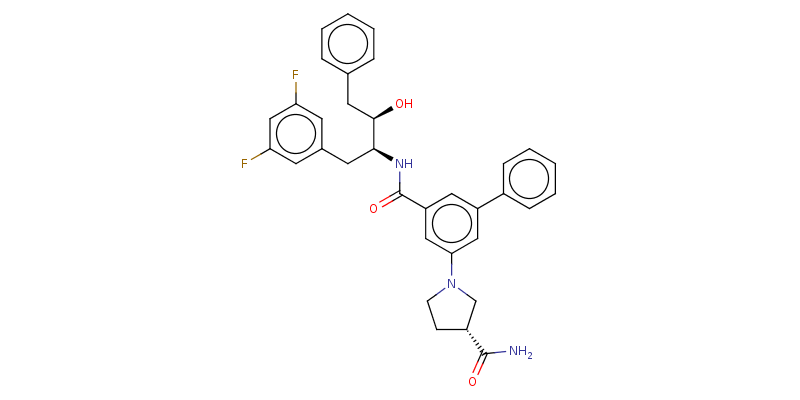 | | |
| C49 | 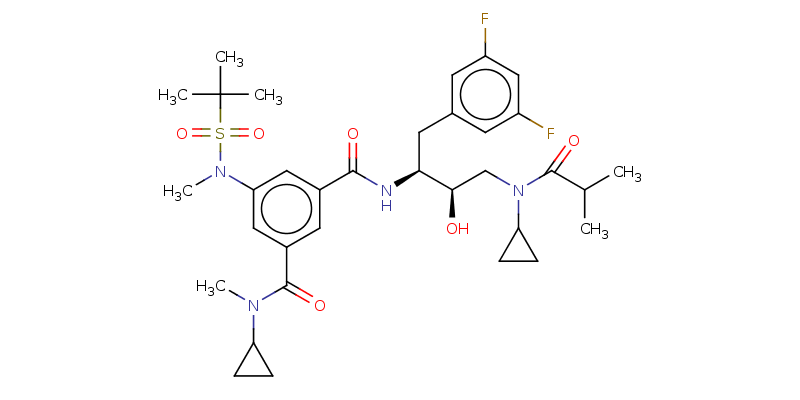 | | |
| C50 | 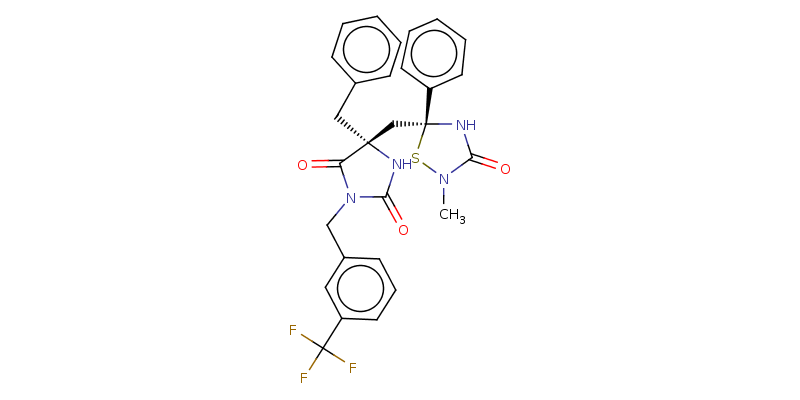 | | |
| C51 | 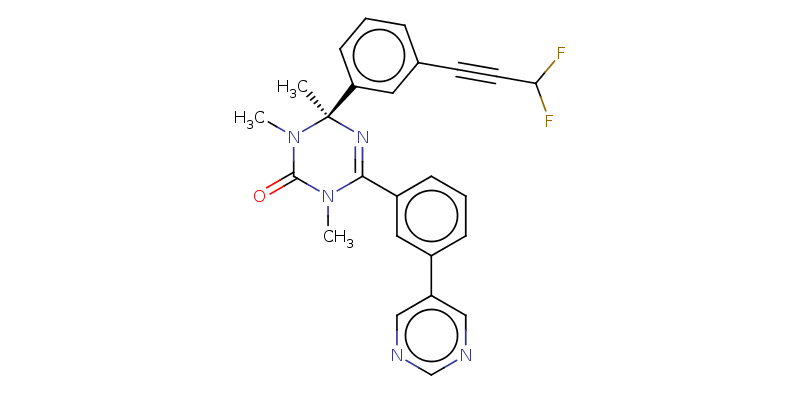 | | |
| C52 | 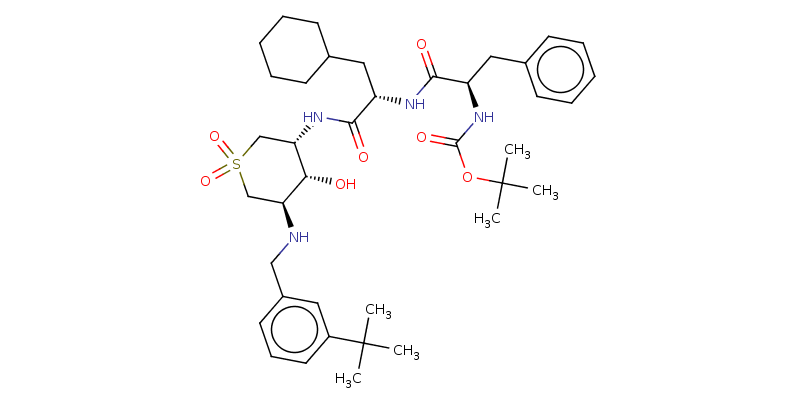 | | |
| C53 | 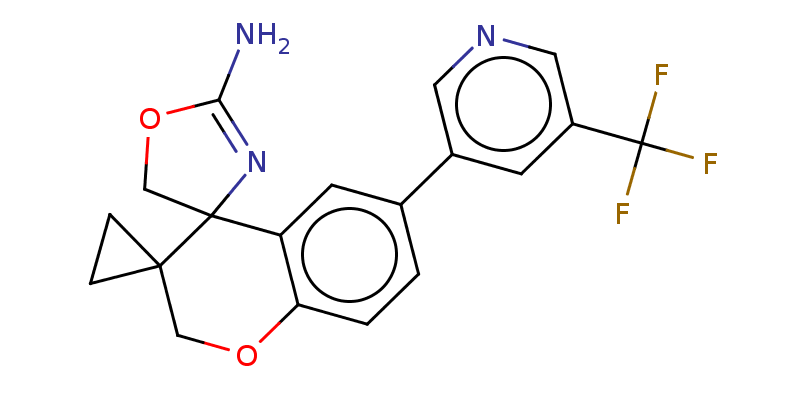 | | |
| C54 | 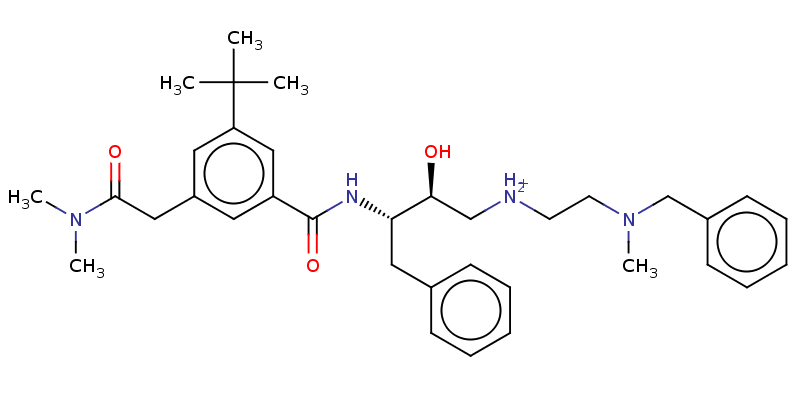 | | |
| C55 | 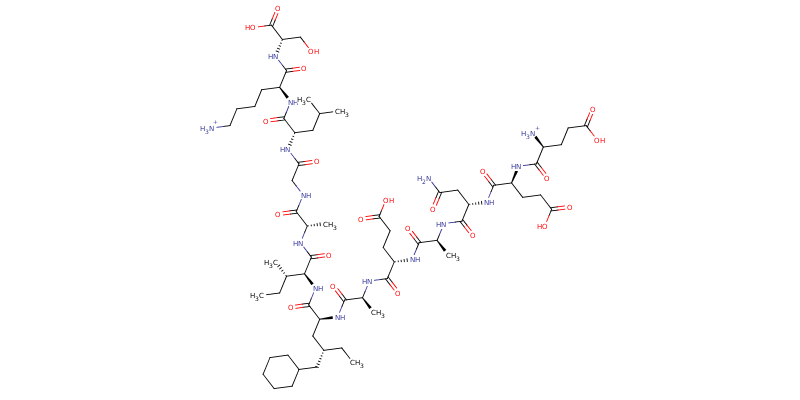 | | |
| C56 | 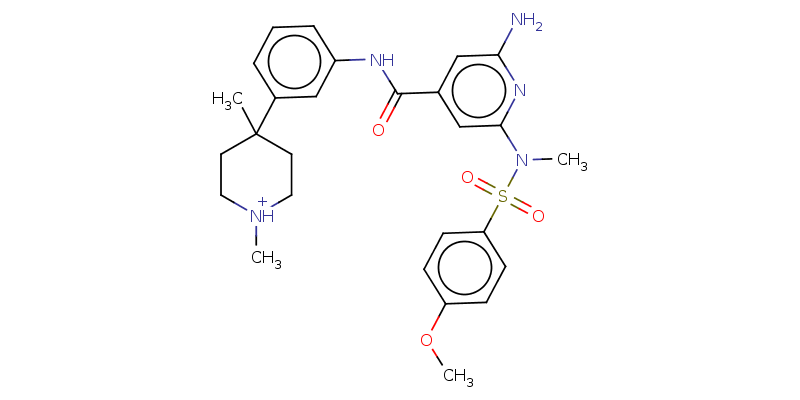 | | |
| C57 | 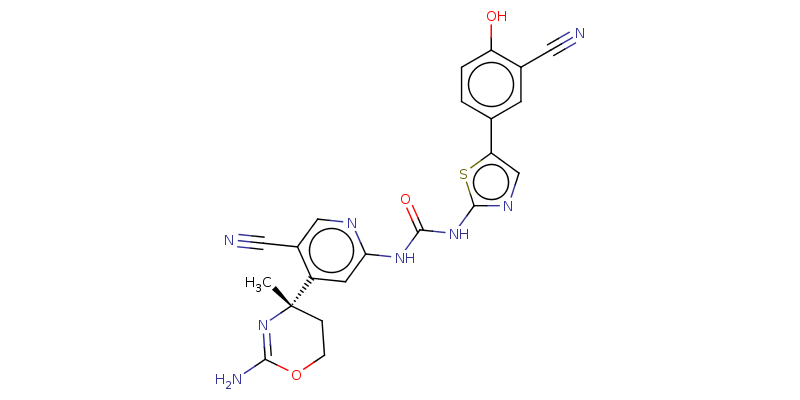 | | |
| C58 | 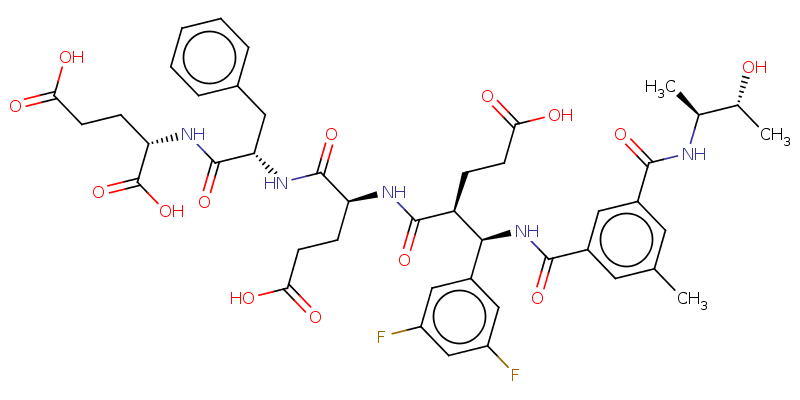 | | |
| C59 | 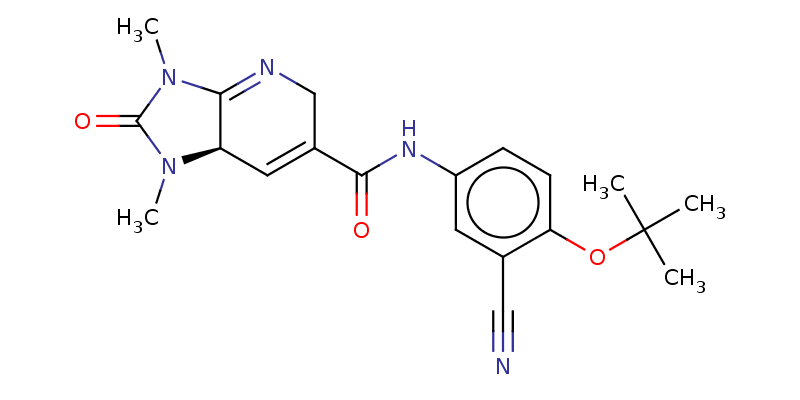 | | |
| C60 | 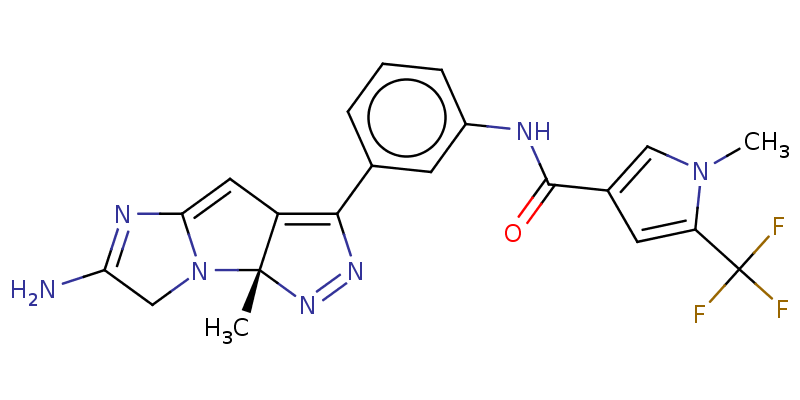 | | |
| C61 | 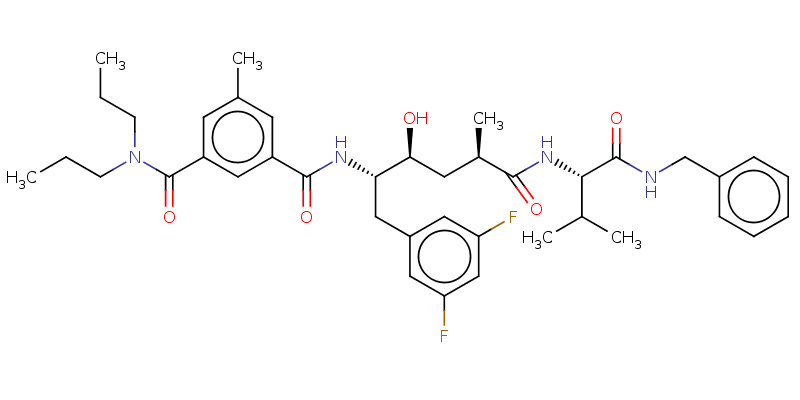 | | |
| C62 | 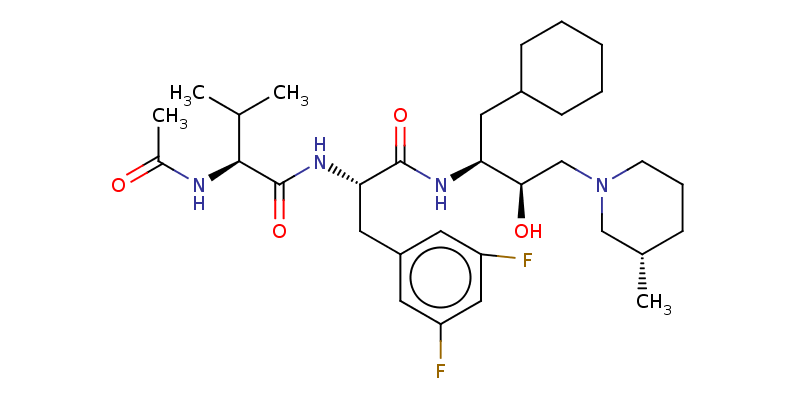 | | |
| C63 | 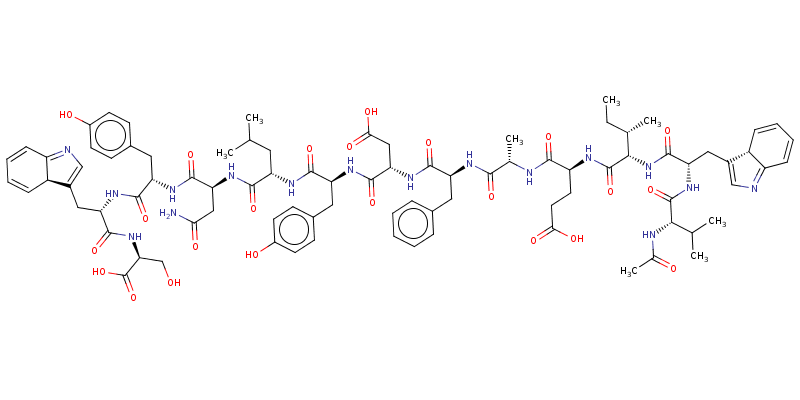 | | |

Table S2: Lipinski RO5 for the 63 compounds generated via DrugGPT

| Compound | Lipinski | | | |
| --- | --- | --- | --- | --- |
|  | R1 | R2 | R3 | R4 |
| C1 | 746.70 | 3.79 | 6 | 7 |
| C2 | 463.61 | 1.83 | 5 | 5 |
| C3 | 693.71 | 3.28 | 5 | 7 |
| C4 | 504.56 | 2.19 | 4 | 6 |
| C5 | 754.72 | 5.65 | 5 | 4 |
| C6 | 449.64 | 1.77 | 3 | 5 |
| C7 | 570.71 | 2.45 | 5 | 5 |
| C8 | 357.41 | 2.52 | 3 | 3 |
| C9 | 747.77 | 0.55 | 7 | 8 |
| C10 | 736.80 | 5.06 | 3 | 6 |
| C11 | 588.64 | 1.70 | 6 | 5 |
| C12 | 558.67 | 3.10 | 4 | 5 |
| C13 | 945.17 | 6.27 | 7 | 9 |
| C14 | 1155.39 | -0.18 | 14 | 14 |
| C15 | 682.91 | 4.98 | 5 | 7 |
| C16 | 458.58 | 1.84 | 3 | 8 |
| C17 | 409.43 | 2.53 | 0 | 5 |
| C18 | 799.87 | 2.73 | 7 | 8 |
| C19 | 1406.58 | 0.68 | 13 | 17 |
| C20 | 415.43 | 2.12 | 2 | 4 |
| C21 | 620.67 | 4.84 | 2 | 5 |
| C22 | 579.78 | 7.64 | 1 | 3 |
| C23 | 583.65 | 4.80 | 3 | 4 |
| C24 | 457.53 | 3.88 | 2 | 6 |
| C25 | 475.59 | 5.34 | 2 | 4 |
| C26 | 560.61 | 2.66 | 5 | 7 |
| C27 | 487.02 | 5.73 | 5 | 3 |
| C28 | 617.75 | 4.97 | 3 | 6 |
| C29 | 446.53 | 4.25 | 2 | 3 |
| C30 | 479.60 | 3.88 | 3 | 3 |
| C31 | 477.51 | 1.63 | 3 | 8 |
| C32 | 676.84 | 5.59 | 5 | 5 |
| C33 | 444.51 | 3.22 | 2 | 5 |
| C34 | 434.41 | 3.85 | 3 | 4 |
| C35 | 826.04 | 4.30 | 6 | 8 |
| C36 | 602.73 | 2.99 | 5 | 6 |
| C37 | 691.89 | 3.70 | 5 | 7 |
| C38 | 476.08 | 4.14 | 4 | 3 |
| C39 | 553.63 | 4.78 | 0 | 7 |
| C40 | 489.56 | 5.24 | 2 | 3 |
| C41 | 863.11 | 5.18 | 6 | 7 |
| C42 | 559.64 | 3.21 | 4 | 5 |
| C43 | 693.84 | 3.41 | 3 | 8 |
| C44 | 432.42 | 2.40 | 3 | 5 |
| C45 | 314.37 | 3.75 | 1 | 2 |
| C46 | 582.75 | 3.94 | 3 | 3 |
| C47 | 488.57 | 3.25 | 3 | 7 |
| C48 | 506.69 | -2.17 | 5 | 4 |
| C49 | 569.65 | 4.88 | 3 | 4 |
| C50 | 676.82 | 4.11 | 2 | 6 |
| C51 | 554.59 | 5.28 | 2 | 4 |
| C52 | 445.47 | 4.37 | 0 | 4 |
| C53 | 726.98 | 4.30 | 5 | 8 |
| C54 | 375.34 | 3.48 | 1 | 5 |
| C55 | 573.80 | 3.01 | 3 | 4 |
| C56 | 1442.67 | -5.45 | 20 | 18 |
| C57 | 524.66 | 2.31 | 3 | 6 |
| C58 | 474.50 | 3.24 | 4 | 10 |
| C59 | 911.90 | 2.23 | 10 | 10 |
| C60 | 381.43 | 2.37 | 1 | 5 |
| C61 | 441.41 | 3.71 | 2 | 7 |
| C62 | 706.87 | 5.72 | 4 | 5 |
| C63 | 592.771 | 3.70 | 4 | 5 |

R1 to R4 denote the Lipinski’s RO5 where R1 = Molecular Weight, R2 = Number of Hydrogen bond donors, R3 = Number of Hydrogen bond acceptors and R4 = LogP

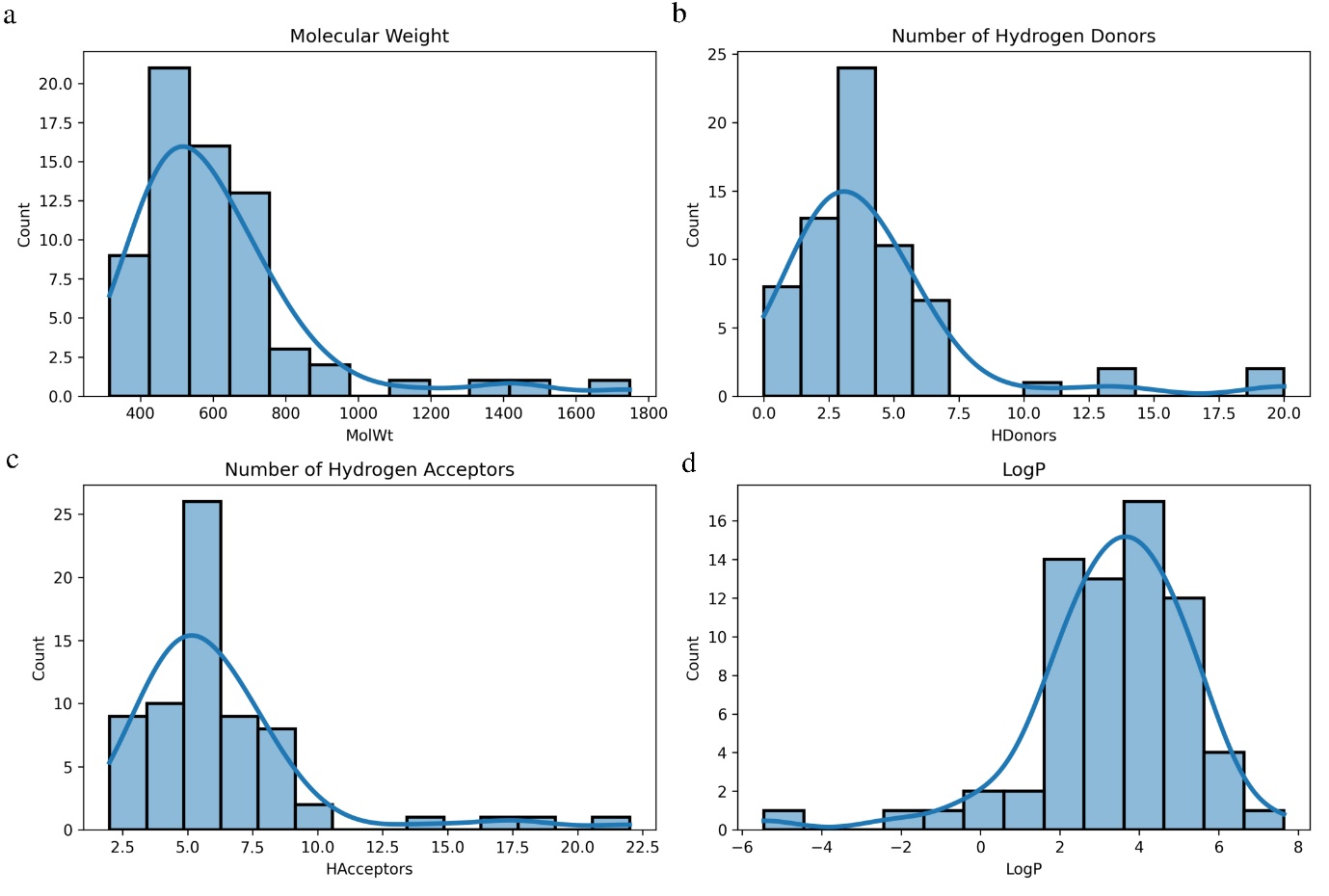

**Figure S1: Distribution analysis of Lipinski’s RO5 for the generated molecules**. The plot represents the distribution of (a) Molecular Weight (b) Number of Hydrogen bond Donors (c) Number of Hydrogen bond acceptors (d) LogP

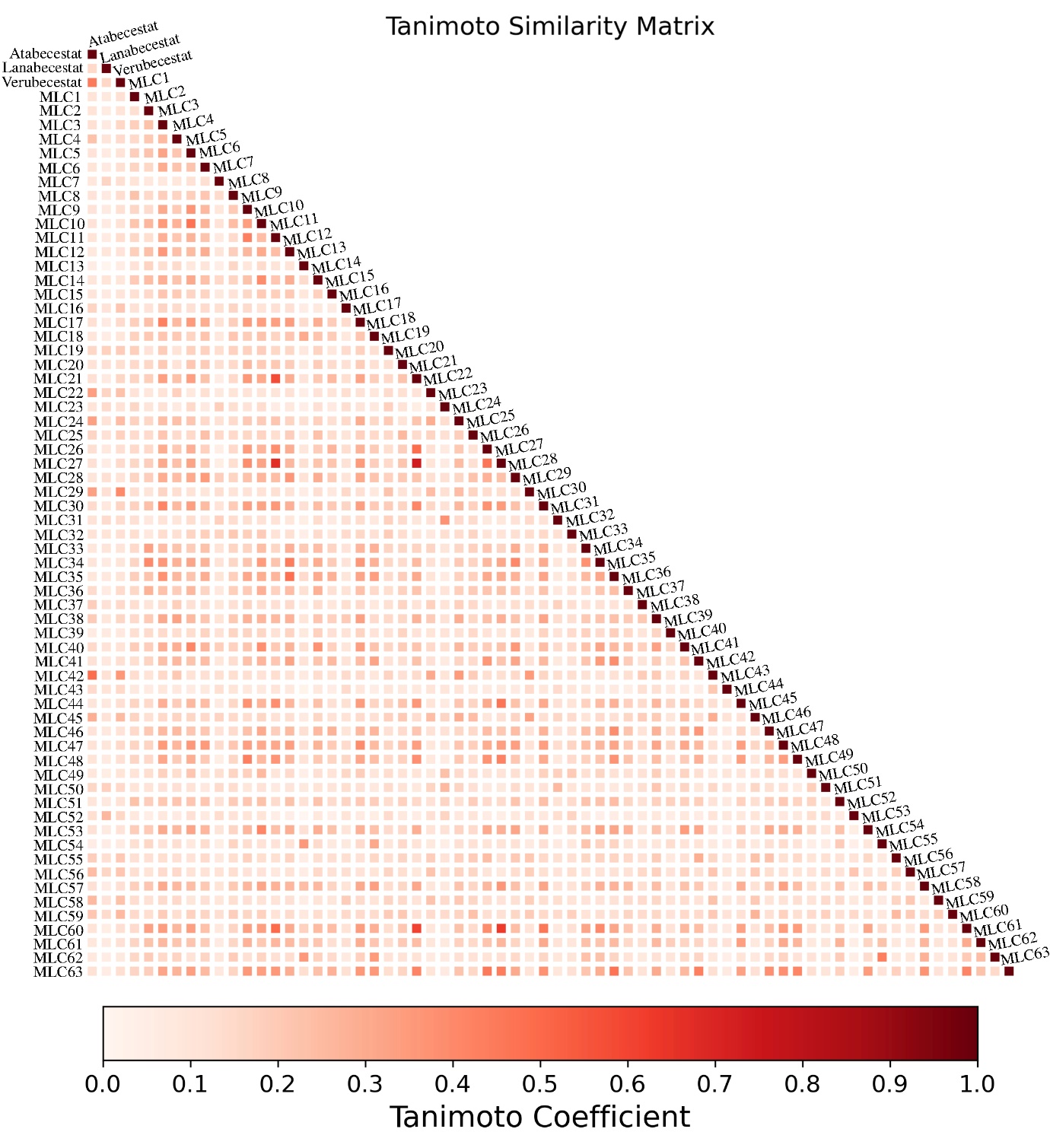

**Figure S2: Tanimoto Similarity of the generated ligands from DrugGPT**. The colorbar represents the Tanimoto coefficient reflecting how similar two compounds are

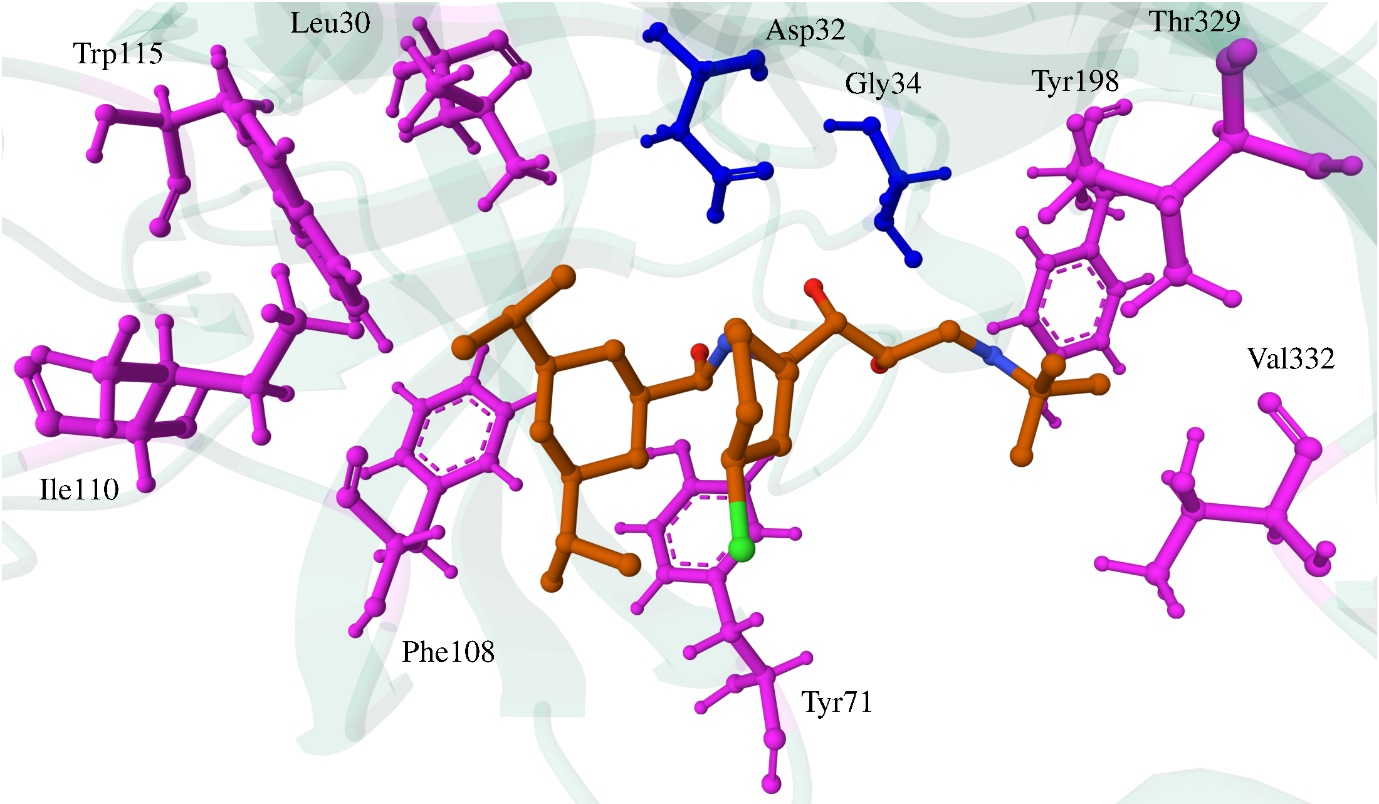

Fig S3: Molecular Docking interaction of MLC11 with BACE1 residues. Blue colour residues form hydrogen bonding and magenta colour residues form hydrophobic interactions

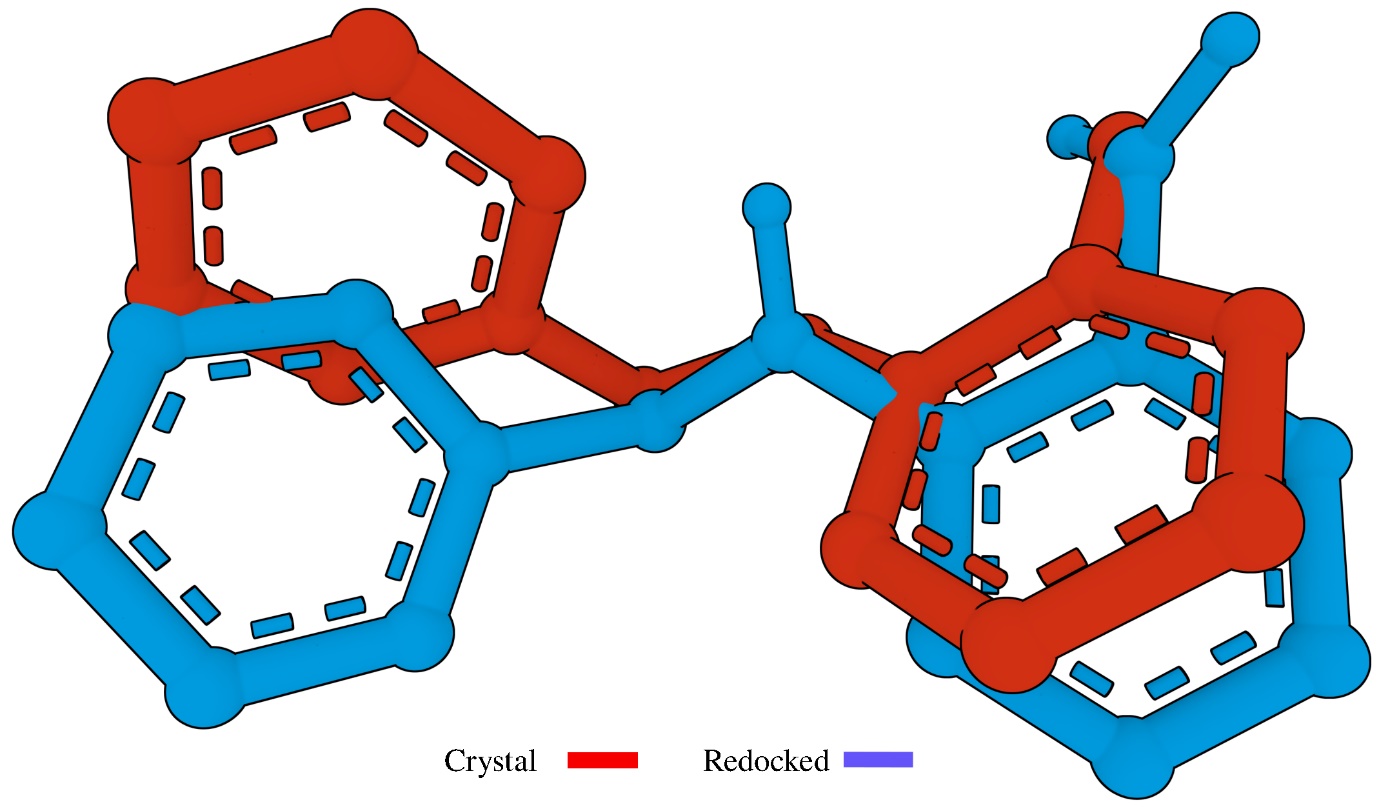

Fig S4: Redocking of 8AP with BACE1. The crystal structure of 8AP (red) is overlapped with the re-docked 8AP (blue) giving an RMSD of 1.1 Å

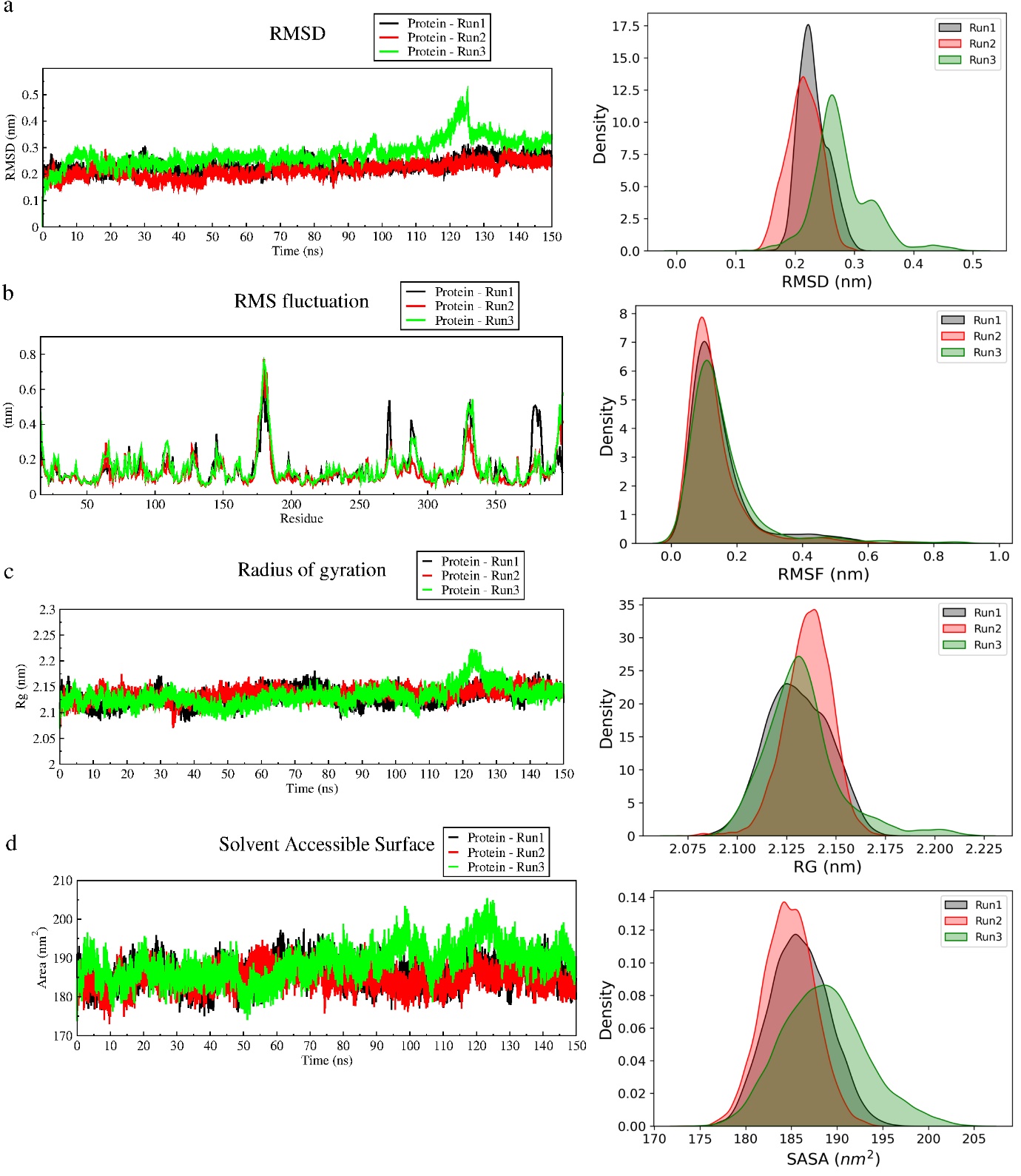

**Fig S5: Triplicate 150 ns MD Analysis of apo BACE1 with run 1, run 2 and run 3 depicted in black, red and green colour respectively**. (a) RMSD (b) RMSF (c) Radius of Gyration (d) SASA. The corresponding probability distribution of the respective properties are provided with the same colour code.

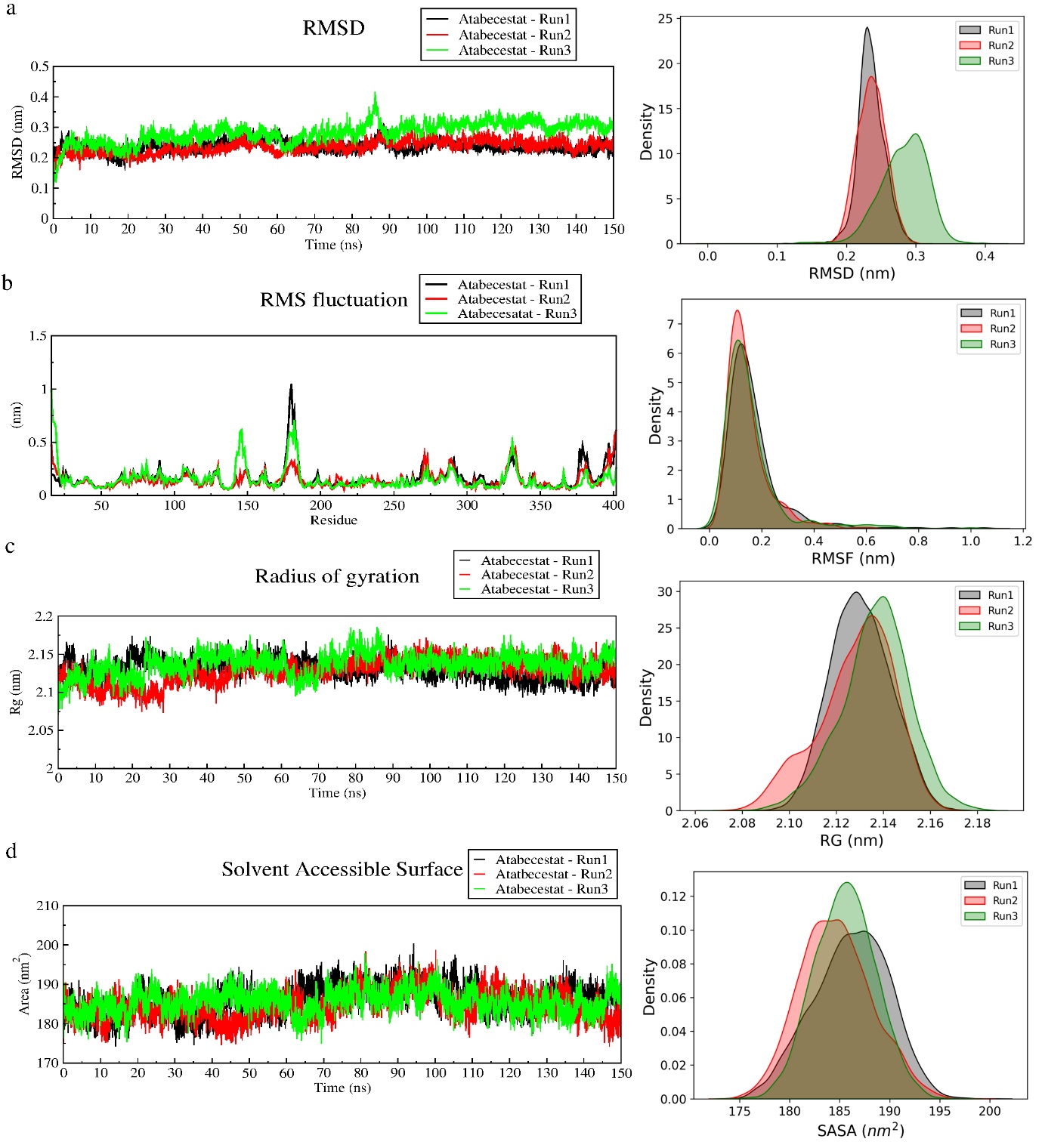

**Fig S6: Triplicate 150 ns MD Analysis of BACE1 with Atabecestat with run 1, run 2 and run 3 depicted in black, red and green colour respectively**. (a) RMSD (b) RMSF (c) Radius of Gyration (d) SASA. The corresponding probability distribution of the respective properties are provided with the same colour code.

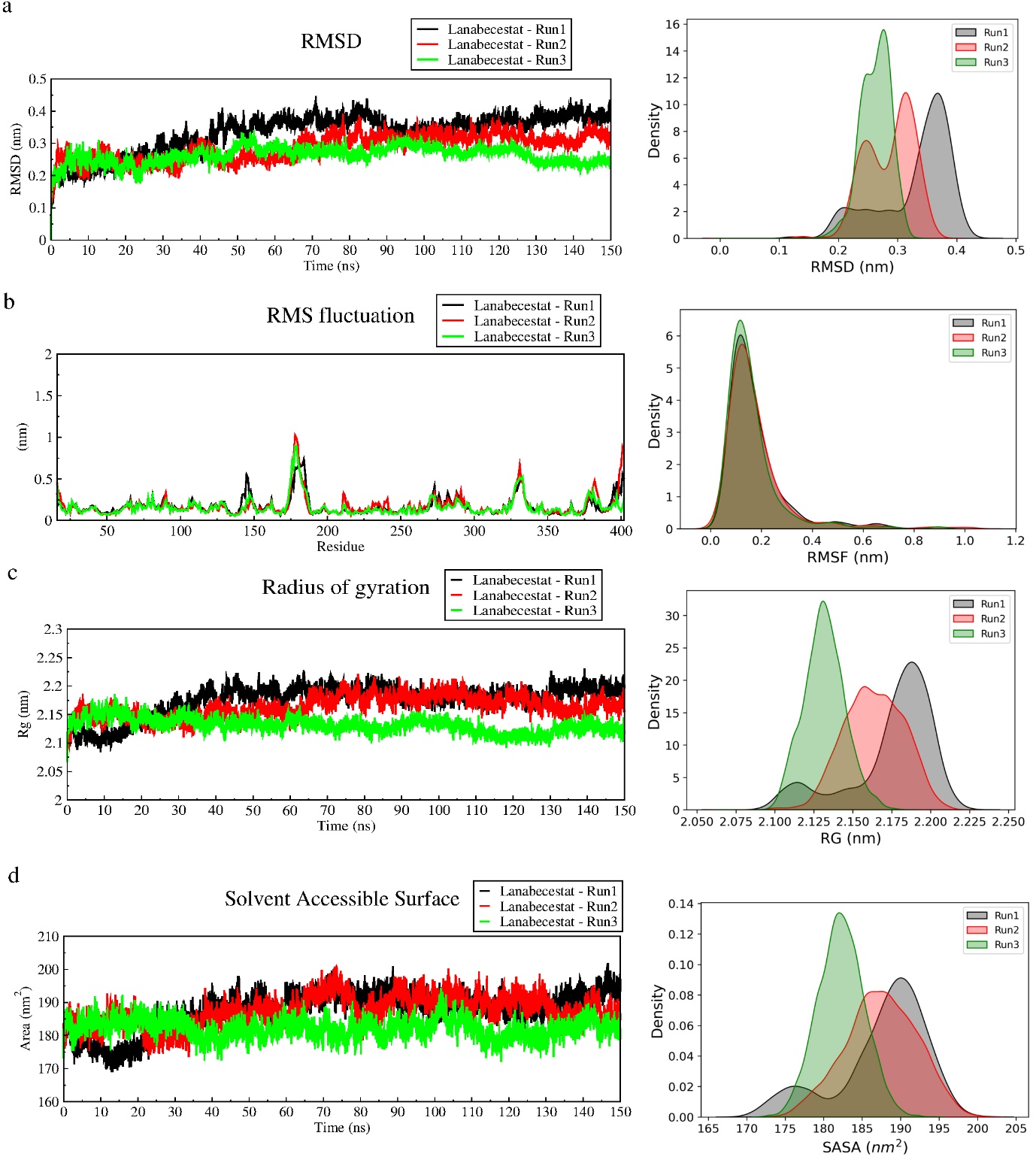

**Fig S7: Triplicate 150 ns MD Analysis of BACE1 with Lanabecestat with run 1, run 2 and run 3 depicted in black, red and green colour respectively**. (a) RMSD (b) RMSF (c) Radius of Gyration (d) SASA. The corresponding probability distribution of the respective properties are provided with the same colour code.

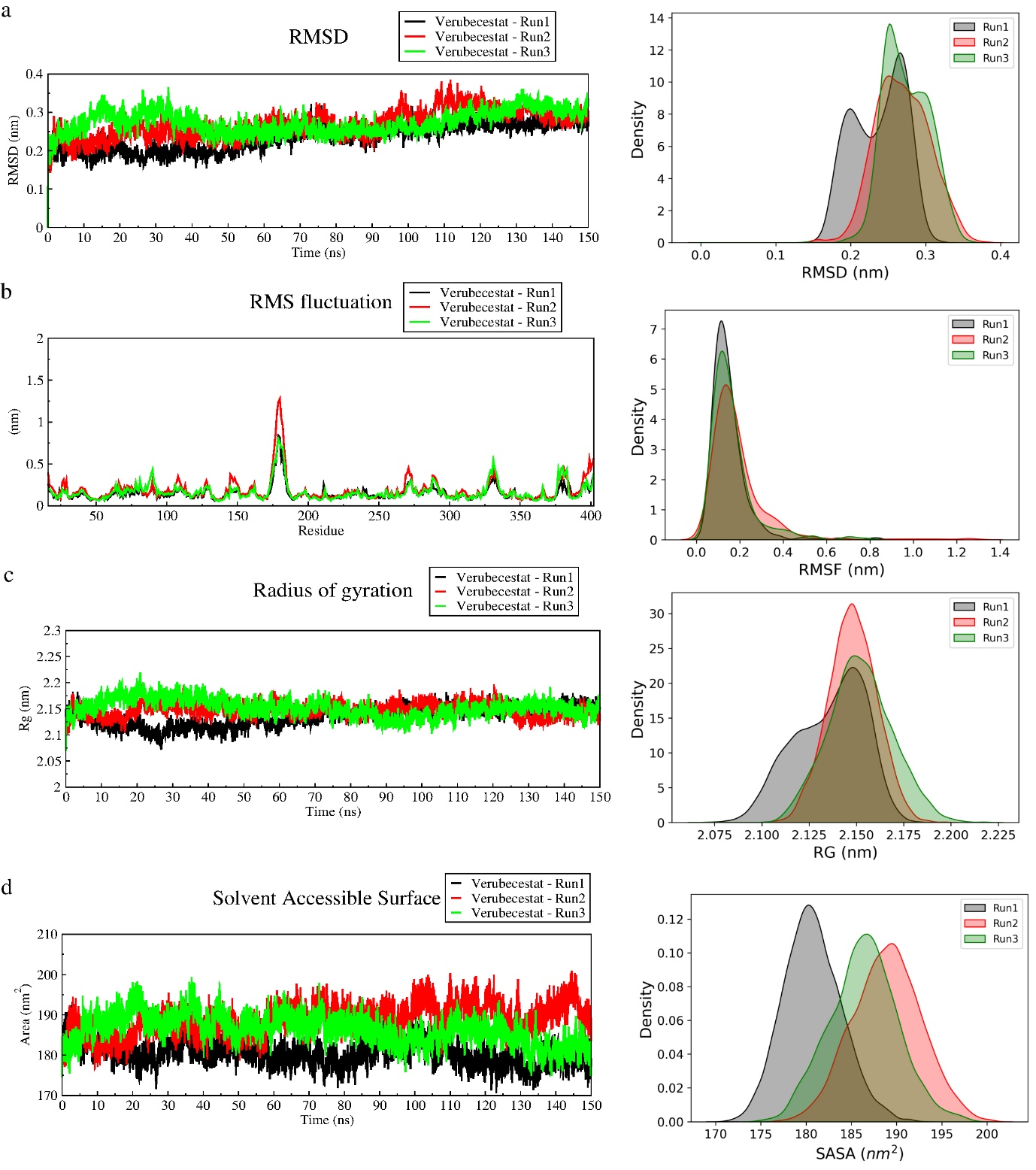

**Fig S8: Triplicate 150 ns MD Analysis of BACE1 with Verubecestat with run 1, run 2 and run 3 depicted in black, red and green colour respectively**. (a) RMSD (b) RMSF (c) Radius of Gyration (d) SASA. The corresponding probability distribution of the respective properties are provided with the same colour code.

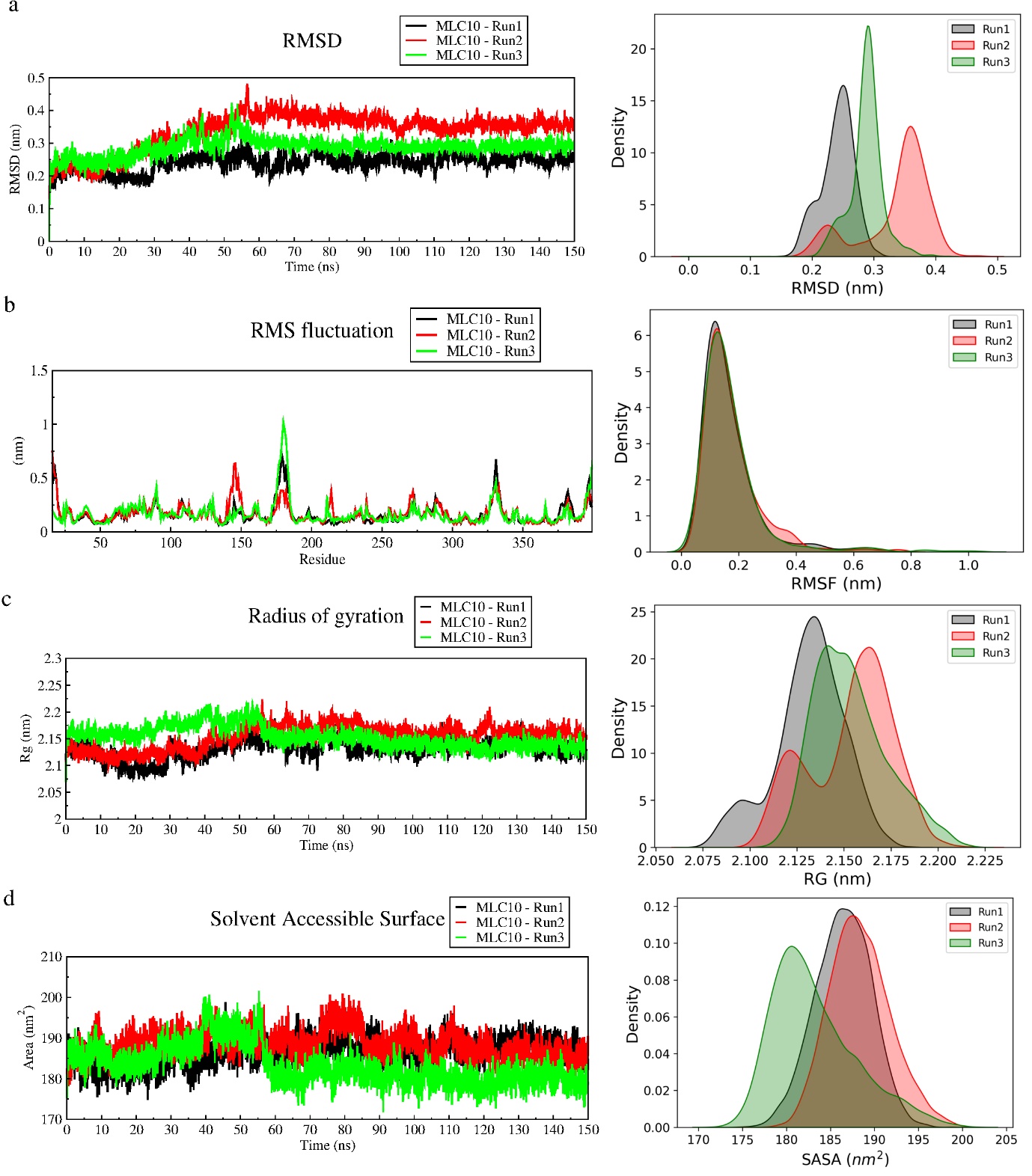

**Fig S9: Triplicate 150 ns MD Analysis of BACE1 with MLC10 with run 1, run 2 and run 3 depicted in black, red and green colour respectively**. (a) RMSD (b) RMSF (c) Radius of Gyration (d) SASA. The corresponding probability distribution of the respective properties are provided with the same colour code.

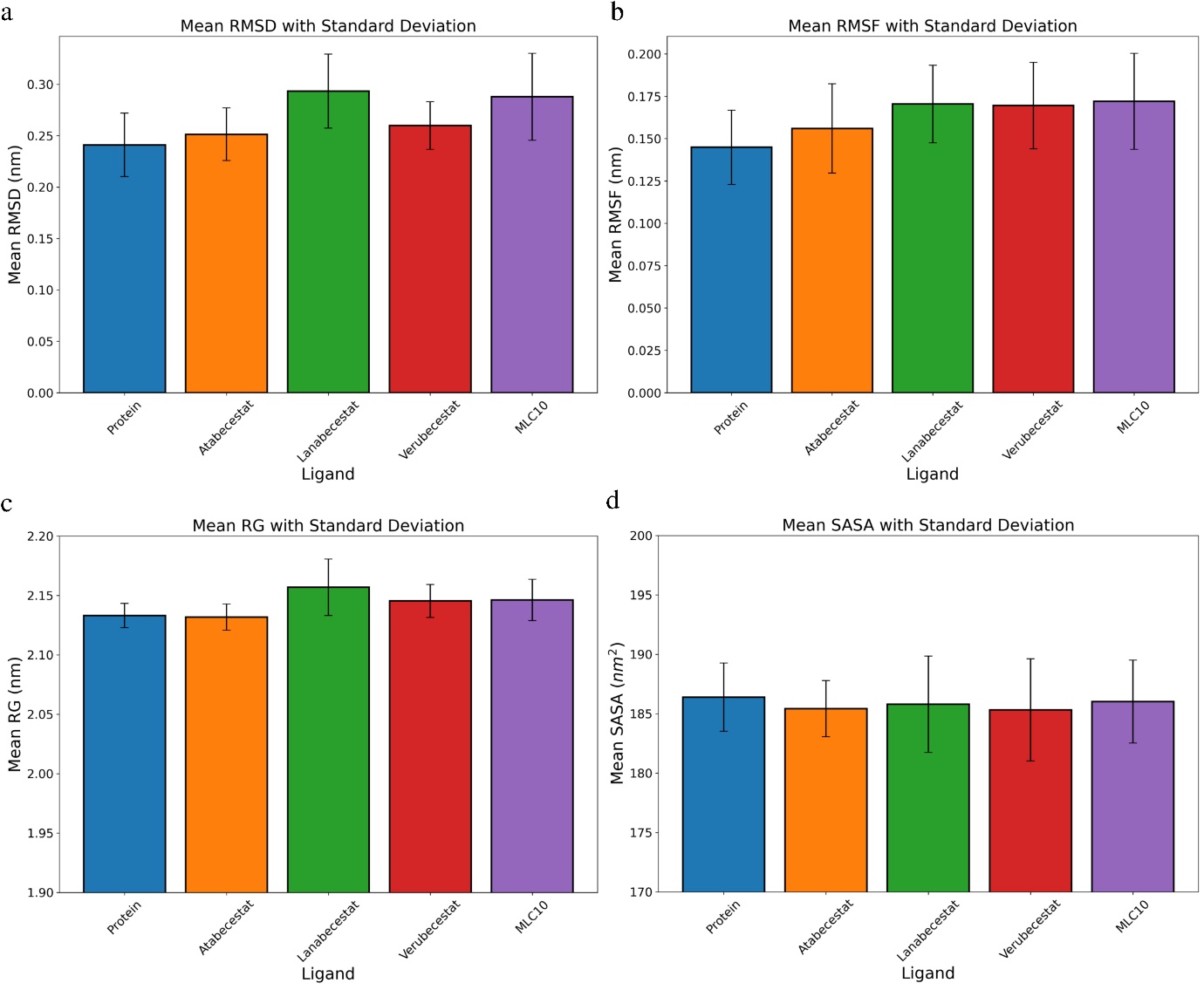

**Fig S10: 150 ns triplicate MD Simulation analysis**. (a) RMSD (b) RMSF (c) Radius of Gyration (d) SASA. The error bar corresponds to the standard deviation associated with each ligand

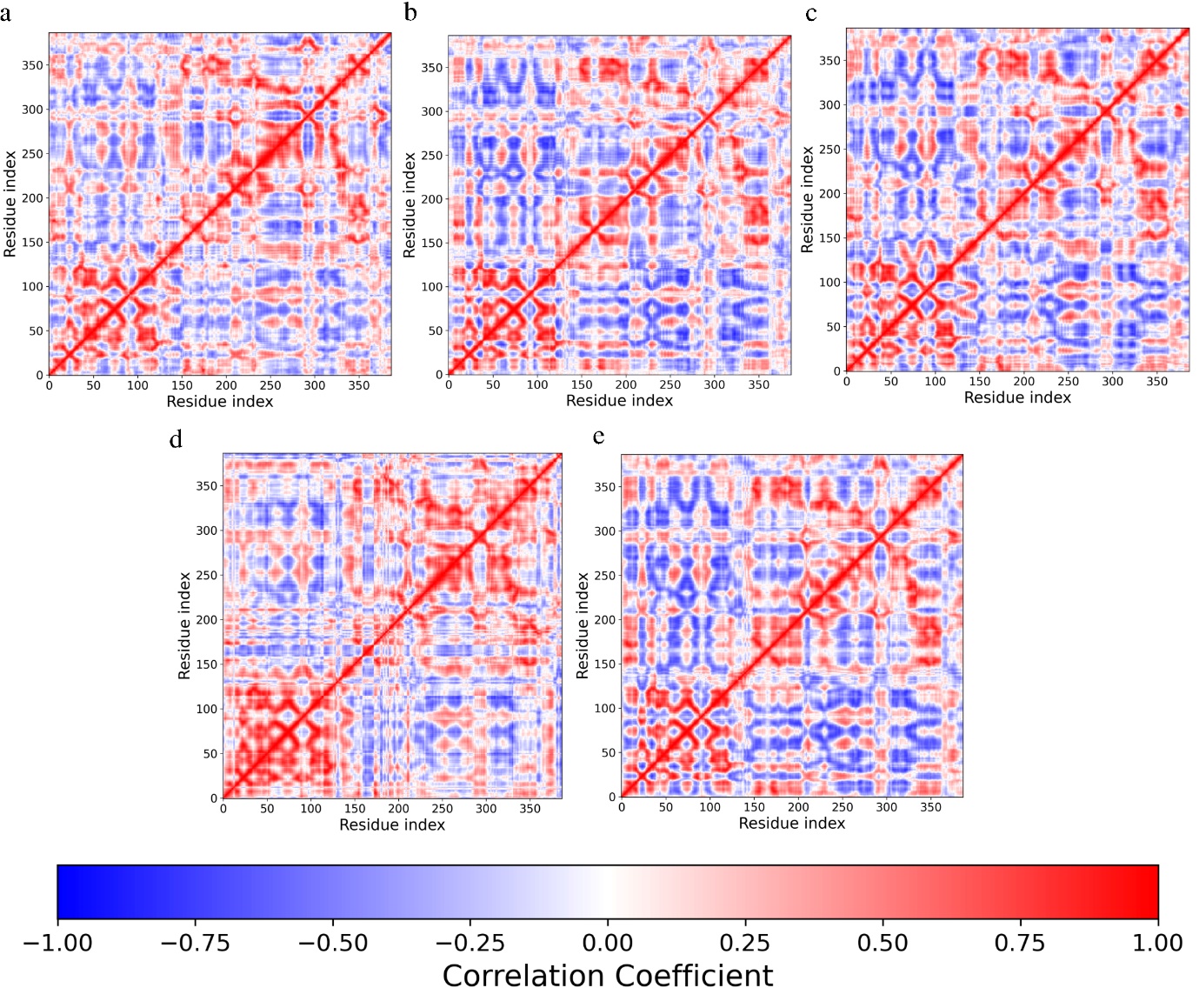

**Fig S11: Dynamic Cross Correlation Matrix (DCCM) analysis for BACE1.** (a) Apo form (b) with Atabecestat (c) with Lanabecestat (d) with Verubecestat (e) with MLC10. The results are averaged correlation of triplicate runs

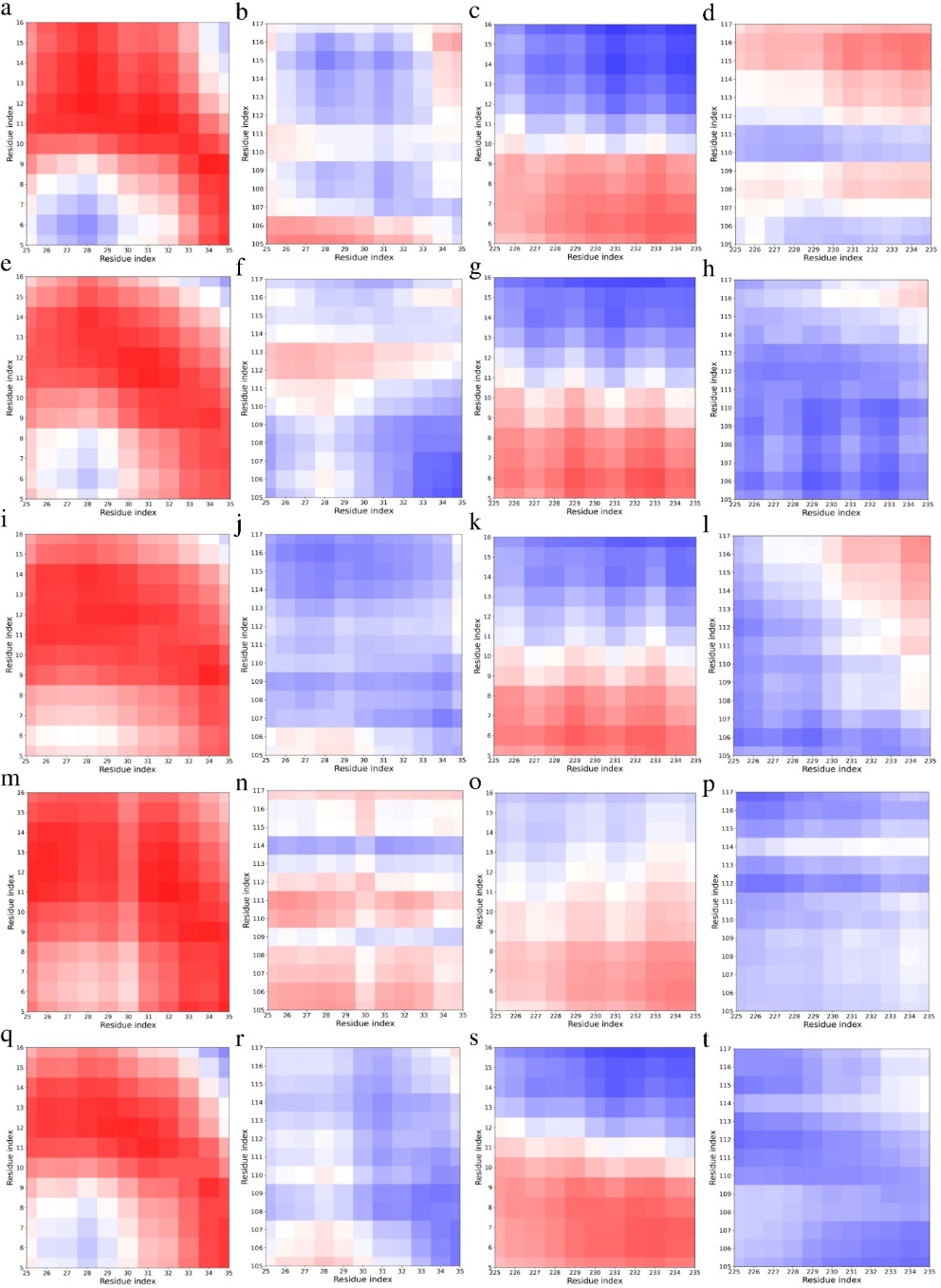

**Fig S12: DCCM analysis for the 150 ns MD simulation of Apo form of BACE1** **with known inhibitors and MLC10.** BACE1 Apo (first row), in association with Atabecestat (second row), in association with Lanabecestat (third row), in association with Verubecestat (forth row), in association with MLC10 (fifth row). The first column corresponds to DCCM between active site 1 (includes Asp32) and 10s loop, the second column corresponds to DCCM between active site 1 and 113s loop, the third column corresponds to DCCM between active site 2 and 10s loop, the fourth column corresponds to DCCM between active site 2 and 113s loop. The colour bar denotes the correlation coefficient with blue corresponding to anti-correlation and red corresponding to positive correlation. The results are averaged correlation of triplicate runs

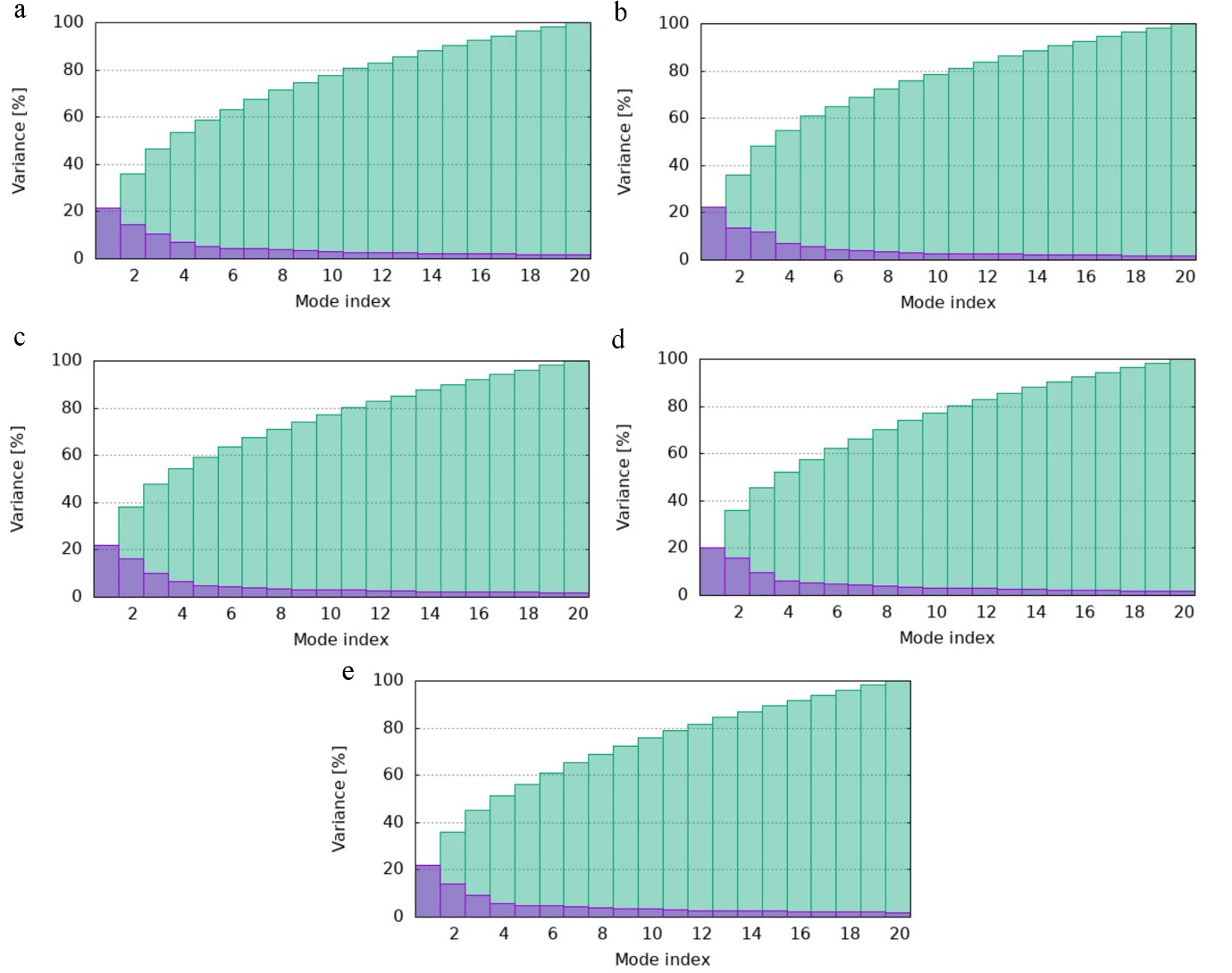

**Fig S13: Variance vs Mode Index for BACE1 NMA Analysis.** (a) Apo form (b) with Atabecestat (c) with Lanabecestat (d) with Verubecestat (e) with MLC10

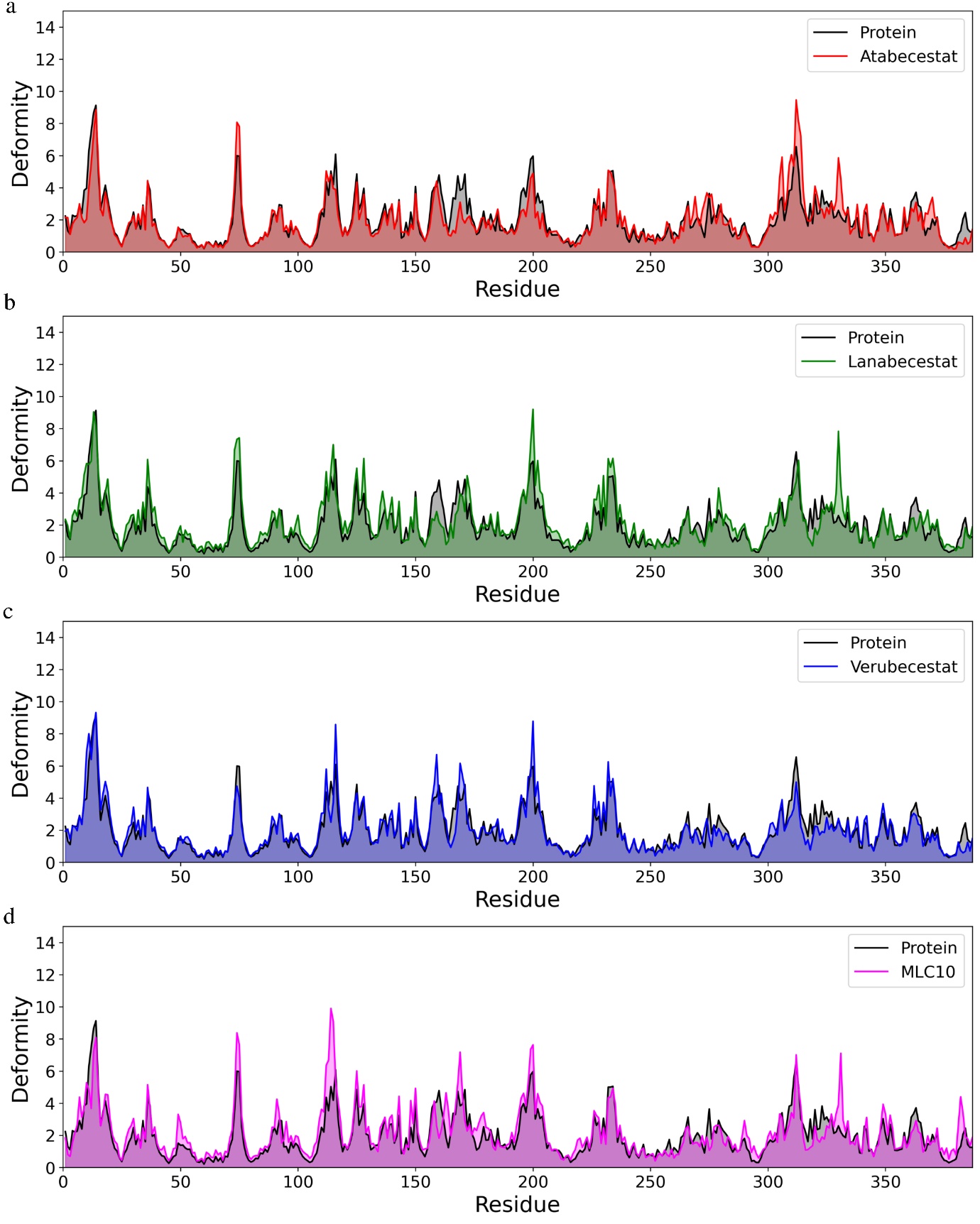

**Fig S14: Deformity analysis with respect to residue position of BACE1.** (a) with Atabecestat vs apo form (b) with Lanabecestat vs apo form (c) with Verubecestat vs apo form (a) with MLC10 vs apo form

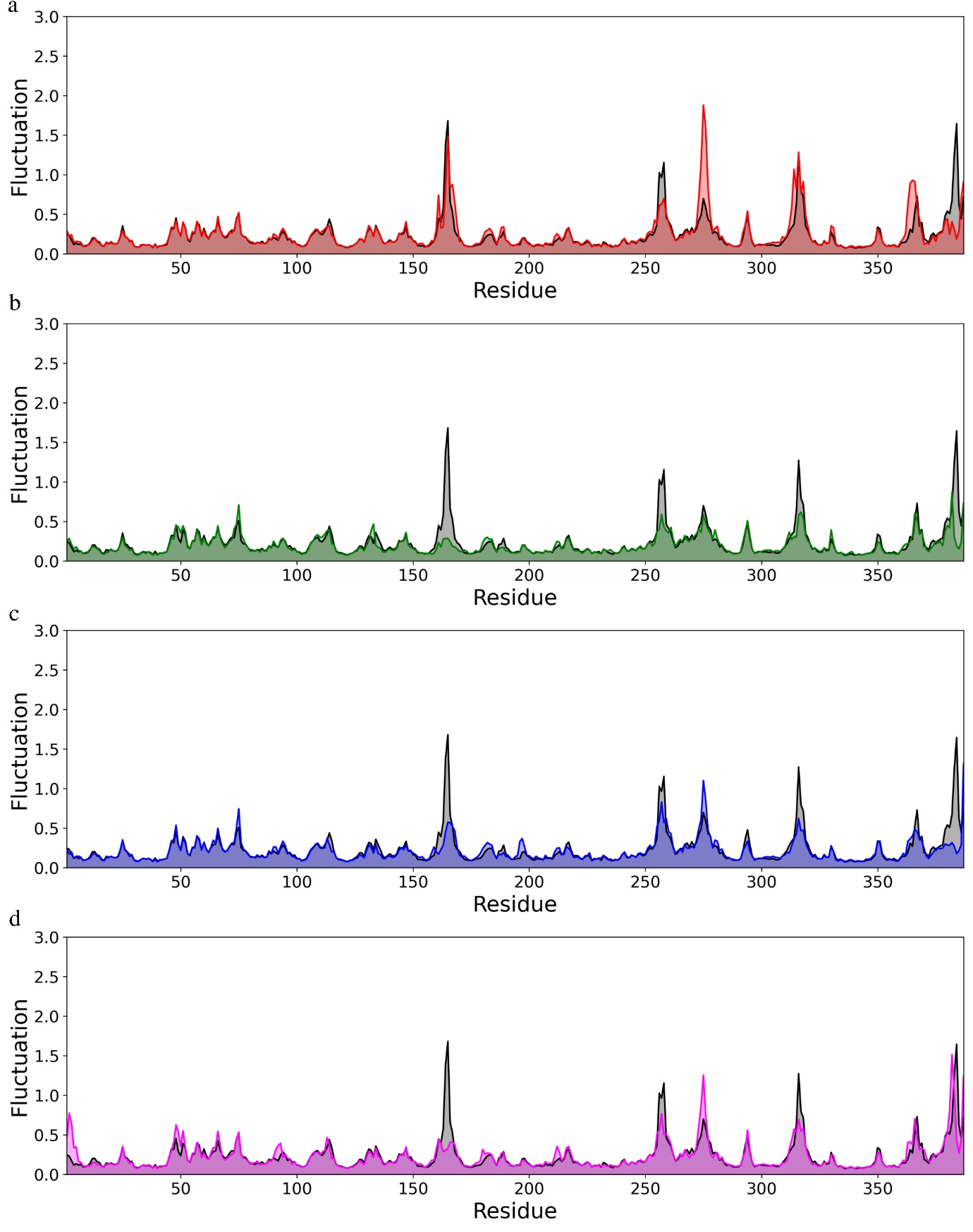

**Fig S15: NMA based fluctuation analysis with respect to residue position of BACE1.** (a) with Atabecestat vs apo form (b) with Lanabecestat vs apo form (c) with Verubecestat vs apo form (a) with MLC10 vs apo form
